# Student-Driven Microbiome Exploration: A Low-Cost 16S rRNA Sequencing Curriculum for Undergraduate Biology Education

**DOI:** 10.64898/2026.05.11.724446

**Authors:** Hassan Barakat, Jingjing Cheng, Makenzie Bolton, Kyle Lee, Ana Vindas, Chance Stephens, Jose Salazar Guerreiro, Ackshaya Muthu Saravanan, Xiangpeng Li

## Abstract

Microbiome science is increasingly important in modern biology education because microbial communities influence human health, ecosystems, and environmental processes. However, undergraduate microbiome instruction is often limited by the high cost and technical complexity of sequencing-based workflows, restricting opportunities for authentic student-driven research. To address this challenge, we developed a low-cost, inquiry-based curriculum that enables undergraduate students to conduct complete microbiome studies using 16S rRNA gene sequencing.

The module integrates project design, environmental sample collection, microbial cell processing, PCR amplification, sequencing, and bioinformatic analysis using open-source tools such as QIIME 2. Cost-reduction strategies included centrifugation-based cell collection and a surfactant-assisted direct PCR workflow that eliminated the need for commercial DNA extraction kits. Students designed independent research projects investigating microbial communities in local environments, including campus water sources and gym equipment surfaces.

Assessment data from post-course surveys, knowledge checks, and student research products demonstrated strong learning gains in microbiome concepts, molecular biology techniques, scientific communication, and computational analysis. Students reported high confidence in PCR, experimental design, and microbiome interpretation, while also identifying bioinformatics as the most challenging yet rewarding component of the curriculum. All participants expressed increased interest in future research in microbiology or bioinformatics.

Overall, this curriculum provides an accessible, scalable framework for integrating next-generation sequencing into undergraduate education while promoting inquiry-driven learning, student ownership, and engagement in authentic scientific research.

## Introduction

Microbiome science has become an essential component of modern biology, transforming our understanding of health, disease, and ecosystems^[1],[2]^. Microorganisms influence nearly every aspect of life on Earth—from human physiology to environmental nutrient cycling—and studying these complex communities offers students a powerful lens through which to explore biological diversity^[2]–[6]^. As microbiome research continues to expand, the ability to design, execute, and interpret microbial community analyses is increasingly valuable for students pursuing careers in biology, medicine, and environmental science^[7]–[10]^.

Despite its relevance, hands-on microbiome education remains limited in many undergraduate programs. Implementing genetic sequencing-based projects often poses significant challenges due to high costs, complex workflows, and the need for bioinformatics expertise^[11]–[13]^. As a result, microbiome-related concepts are commonly taught through demonstrations or pre-analyzed datasets rather than authentic, student-driven investigations^[14],[15]^. This disconnect can reduce engagement and limit opportunities for students to experience the iterative and exploratory nature of scientific research.

Inquiry-based and student-driven learning approaches have been shown to improve student motivation, deepen conceptual understanding, and enhance scientific reasoning skills^[16]–[19]^. Course-based Undergraduate Research Experiences (CUREs) extend these benefits to a broader student population by embedding authentic research within formal coursework^[20],[21]^. CUREs in microbiology often involve culturing microorganisms or exploring antibiotic resistance, yet few provide opportunities for students to engage directly with DNA sequencing and computational data analysis due to logistical and financial constraints^[15],[22]–[24]^.

To bridge this gap, we developed a low-cost, modular 16S rRNA sequencing curriculum that enables undergraduate students to perform their own microbiome investigations from start to finish. The module guides students through each stage of the research process: developing a research question, collecting samples from self-selected environments, extracting and amplifying microbial DNA, and analyzing sequencing results using open-source bioinformatics tools such as QIIME 2^[25],[26]^. The workflow integrates molecular and computational components in a format suitable for teaching laboratories with limited budgets or resources.

The curriculum is designed to promote three key educational outcomes: (1) enhance student understanding of microbial ecology and sequencing technologies, (2) build confidence and ownership through inquiry-driven project design, and (3) strengthen computational literacy by engaging students in bioinformatic analysis. This framework supports the American Society for Microbiology (ASM) curricular guidelines emphasizing scientific thinking, data analysis, and communication^[27]^.

Here, we describe the design, implementation, and assessment of this student-driven microbiome exploration curriculum. The module provides a scalable and accessible model for incorporating next-generation sequencing into undergraduate biology education while fostering engagement, curiosity, and critical thinking through authentic research experiences.

## Methods

### 1. Curricular Design

This curriculum was developed as part of a Directed Individual Study (DIS) course, providing students with an opportunity to engage in microbiome research within a research laboratory setting. The module is suitable for upper-level undergraduates majoring in microbiology, biology, ecology, chemistry, biochemistry, or bioinformatics, and can be adapted for use in laboratory-based courses emphasizing molecular biology or microbial ecology.

The module can be implemented over four to six laboratory sessions or expanded into a full-semester research experience, depending on institutional scheduling and available resources. In a shorter implementation, students focus on project design, sample collection, sequencing library preparation, sequencing, and data analysis using pre-sequenced datasets. In a semester-long format, students complete the entire workflow— from project design, environmental sampling through sequencing and interpretation—mirroring a full research cycle (**Fig. 1**).

**Figure 1.**
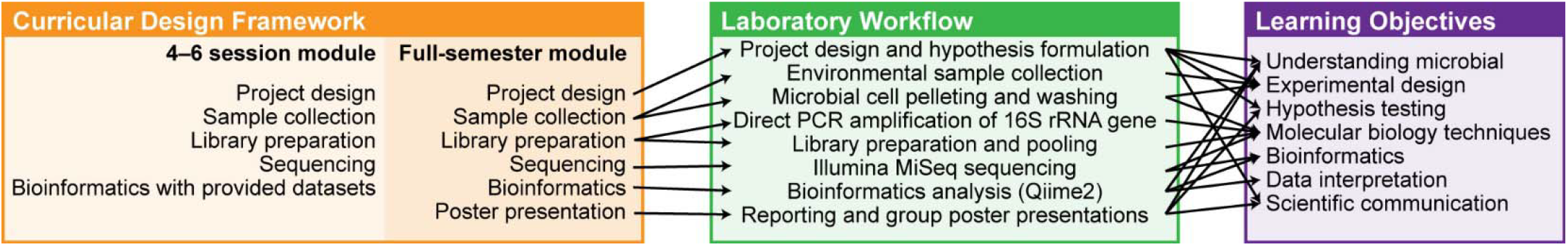
Curricular design framework, laboratory workflow, and learning objectives for the inquiry-based microbiome module. The schematic summarizes the structure and educational alignment of the curriculum. The left panel outlines two implementation formats: a 4–6 session module (project design, sample collection, library preparation, sequencing, and bioinformatics using provided datasets) and a full-semester module that includes the complete research cycle culminating in poster presentations. The central panel depicts the stepwise laboratory workflow, from project design and environmental sampling through microbial cell pelleting and washing, direct PCR amplification of the 16S rRNA gene, library preparation and pooling, Illumina MiSeq sequencing, bioinformatics analysis (QIIME 2), and reporting via group poster presentations. The right panel maps these activities to targeted learning objectives, including understanding microbial communities, experimental design and hypothesis testing, application of molecular biology techniques, bioinformatics analysis, data interpretation, and scientific communication.

To enhance accessibility and affordability, the protocol employs two cost-reduction strategies. First, a centrifugation-based cell collection method is used. Samples collected from water or surfaces (via swabbing into buffer) are centrifuged to pellet microbial cells. The pellet is washed five times with buffer and then transferred to PCR tubes. This approach avoids the higher costs associated with alternative cell isolation techniques such as membrane filtration or commercial extraction kits. Second, the curriculum adopts a direct PCR approach that uses surfactant-assisted lysis and thermal heating during PCR to disrupt cells *in situ*^*[28]*^.

This strategy eliminates the need for separate DNA extraction steps, thereby reducing reagent costs, simplifying the workflow, and decreasing laboratory time while maintaining sufficient template quality for 16S rRNA gene amplification.

Learning objectives were designed to emphasize both conceptual understanding and scientific process skills. Upon completing the module, students are expected to:

- Explain the role of microbial communities in natural and engineered environments and their relevance to human and ecosystem health.
- Design and conduct an independent microbiome study, including hypothesis formulation, sampling strategy, and experimental planning.
- Apply molecular biology techniques such as DNA extraction and 16S rRNA gene amplification to characterize microbial communities^[29]–[31]^.
- Analyze and interpret sequencing data using open-source bioinformatics pipelines (e.g., QIIME 2^[25],[26]^) to evaluate microbial diversity and taxonomy.
- Communicate scientific findings effectively through written reports or poster presentations. This design promotes inquiry-based learning, fosters ownership of research, and integrates molecular and computational skills essential for modern microbiology education. The flexible structure allows adaptation for diverse institutional contexts, supporting both independent study and course-based research experiences.

A complete list of the required laboratory equipment, consumables, and computational resources is provided in the **Supporting Information**. This includes all materials necessary for sample collection, DNA processing, PCR amplification, and sequencing preparation, as well as software tools and hardware requirements for downstream bioinformatic analysis. The SI is intended to ensure full reproducibility of the workflow and to facilitate adoption of the curriculum across teaching and research laboratories with varying levels of infrastructure.

## 2. Laboratory Workflow

The curriculum guides students through each step of a complete 16S rRNA gene sequencing project, from experimental design to data interpretation (**Fig. 2**). The workflow is designed to balance feasibility, cost efficiency, and educational value, allowing students to experience an authentic research process within the constraints of a teaching or research lab.

**Figure 2.**
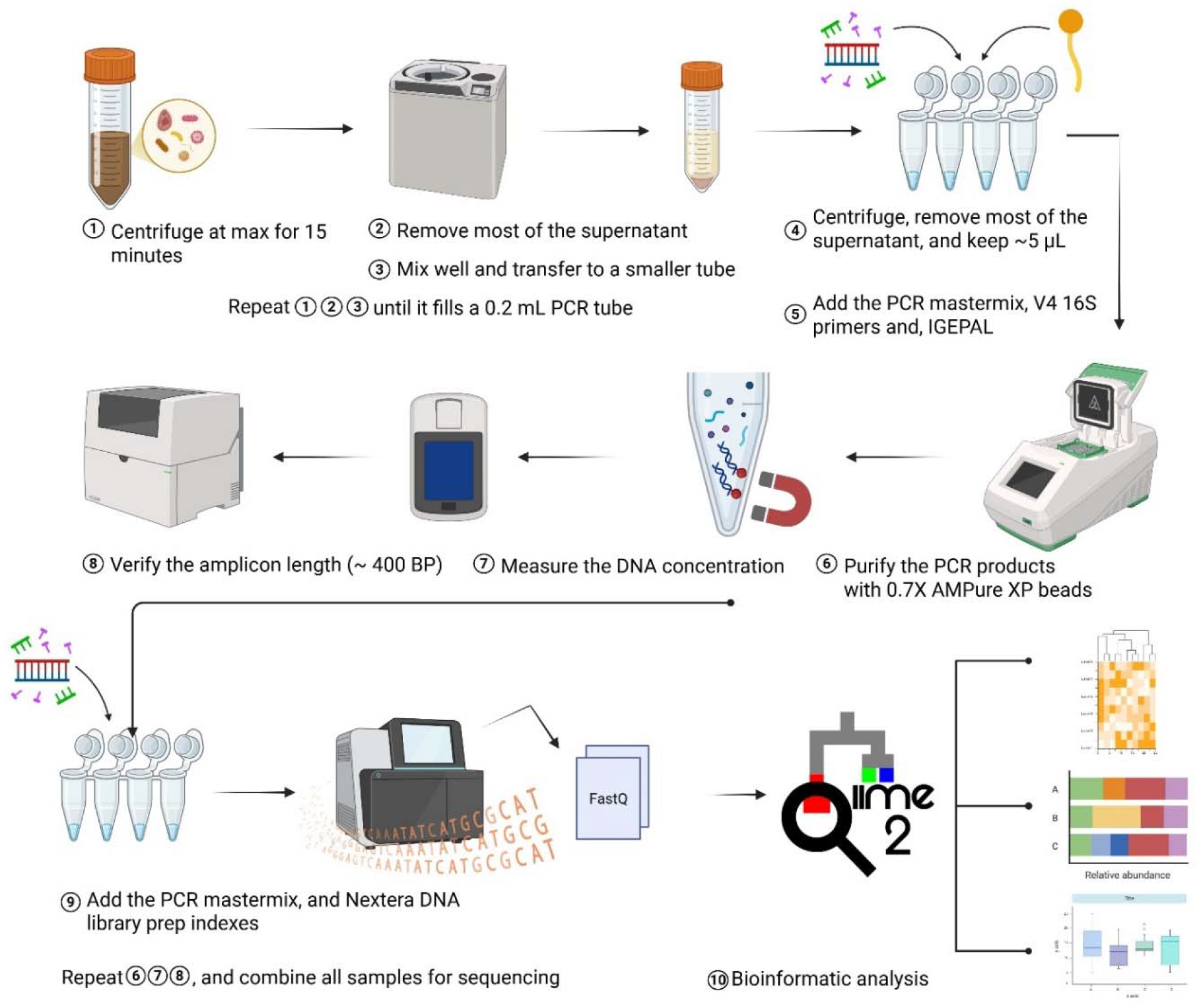
Workflow for 16S rRNA amplicon library preparation and sequencing. Samples were first concentrated by repeated centrifugation and supernatant removal and then transferred into a 0.2 mL PCR tube. After a final centrifugation step, approximately 5 µL of sample was retained and combined with PCR master mix, V4 16S primers, and IGEPAL for amplification. PCR products were purified using 0.7× AMPure XP beads, quantified, and verified for the expected amplicon size (∼400 bp). Indexed sequencing libraries were then generated using Nextera DNA library preparation indexes, followed by additional purification, quantification, and quality control steps prior to pooling and sequencing. Sequencing reads (FASTQ files) were processed using the QIIME 2 pipeline for downstream bioinformatic analyses, including taxonomic profiling and relative

### 2.1. Project Design

The module begins with an introductory lecture on microbial community analysis using 16S rRNA sequencing. In this session, the instructor introduces foundational concepts in microbiome research, including microbial diversity, taxonomic profiling, and the principles of hypothesis-driven investigation. Following the lecture, students design their own microbiome research projects, working individually or in small groups. Each group identifies a research question related to microbial communities in their everyday environments, formulates a testable hypothesis, and develops a sampling and analysis plan.

Throughout this process, faculty and graduate teaching assistants provide structured guidance through class discussions, proposal templates (**Note S1**), and iterative feedback to ensure that each project is scientifically sound, feasible, and aligned with biosafety and ethical standards. Students are required to assess potential risks, obtain appropriate approvals for sampling locations, and implement safe handling practices for environmental materials. This stage emphasizes scientific reasoning, experimental design (including appropriate controls), and the ability to translate broad biological questions into testable hypotheses that can be pursued using a low-cost, hands-on workflow in subsequent lab sessions.

### 2.2. Sample Collection

Students collect samples from self-selected locations such as water or surfaces in built environments (e.g., gym equipment, classroom desks, or library computer keyboards). Standardized sampling materials and biosafety protocols are provided to minimize contamination and ensure reproducibility. For water samples, 50 mL is collected directly into sterile 50 mL conical tubes. For surface sampling, a sterile cotton swab pre-wetted with Tris-HCl buffer containing 0.1% Tween 20 is used to wipe the designated surface and then transferred into a 1.5 mL tube containing the same buffer. The swab is soaked and gently squeezed several times to release bacterial cells into the buffer. After collection, samples are immediately transported to the laboratory and processed in the next step. Each student or team collects multiple replicates to capture microbial diversity within their chosen environment. Detailed process is included in **Fig. 2** and **Note S2**.

### 2.3. Microbial Cell Collection

Microbial cells are pelleted by centrifugation. The cells are washed for 5 times, and transferred into PCR Tube for storage and PCR amplification.

### 2.4. PCR Amplification and Library Preparation

The V3–V4 region of the 16S rRNA gene is amplified using universal primers (515F/806R) with Illumina adapter overhangs^[32]–[34]^. Students prepare reaction mixtures, perform thermocycling, and verify product size and yield by agarose gel electrophoresis (**Fig. S1**). Barcoding and library preparation are completed using a two-step PCR protocol compatible with pooled sequencing^[34]^.

### 2.5. Pooling and Sequencing

Amplicon libraries are pooled and purified before submission for sequencing. Samples are typically outsourced to a commercial sequencing provider using the Illumina MiSeq platform with 2 × 250 bp paired-end reads. Cost per sample is minimized by batching student projects into a single sequencing run, allowing per-sample costs as low as $13 (**Table S1**). Turnaround time and cost-sharing logistics are discussed with students as part of the project management experience.

### 2.6. Bioinformatics and Data Analysis

After sequencing, students perform data analysis using open-source bioinformatics pipelines such as QIIME 2^[25],[26]^, or optionally Python^[35]^ or R^[36]^-based workflows. Students learn to import sequence data, perform quality control, generate amplicon sequence variants (ASVs) or operational taxonomic units (OTUs), and assign taxonomy using reference databases (e.g., SILVA^[37],[38]^ or Greengenes^[39]^). Analyses include calculation of α-and β-diversity metrics, taxonomic composition visualization (bar plots, PCoA), and hypothesis-driven comparisons among samples. A detailed example of the bioinformatic pipeline is given in **Note S3** and **S4**. Emphasis is placed on reproducibility, data interpretation, and the connection between computational results and biological meaning.

### 2.7. Reporting and Scientific Communication

The module culminates in a formal poster presentation in which each group communicates the design, execution, and findings of their microbiome study. This final stage emphasizes scientific communication as an integral component of the research process and reinforces the connection between experimental design, data analysis, and interpretation.

- Each group prepares a research-style poster that includes the following components:
- Introduction and Rationale: Background information, research question, and hypothesis.
- Methods: Sampling strategy, molecular workflow (direct PCR or DNA extraction, 16S rRNA amplification), sequencing approach, and bioinformatics pipeline.
- Results: Key findings presented through figures and tables, such as gel images, diversity metrics (α-and β-diversity), taxonomic bar plots, and ordination analyses (e.g., PCoA).
- Discussion: Interpretation of results in the context of the original hypothesis, potential sources of bias or experimental limitations, and broader biological or environmental implications.
- Future Directions: Proposed follow-up experiments or methodological improvements.

Students are encouraged to present data visually and concisely, applying principles of effective figure design and scientific storytelling. Instructors provide guidelines on poster organization, graphical clarity, and proper citation of references and analytical tools. Draft posters may undergo peer review prior to the final presentation to promote iterative improvement and collaborative learning.

The module concludes with a mini-symposium in which students present their posters to classmates, faculty, and invited guests. During this session, students respond to questions, justify methodological decisions, and reflect on challenges encountered during the project. Assessment is based on scientific rigor, clarity of communication, quality of data interpretation, and the ability to connect computational analyses to biological meaning. The module concludes with a mini-symposium in which students present their posters to classmates, faculty, and invited guests. Students are also encouraged to present their findings at undergraduate research symposia or local chapters of ASM conferences^[40]^.

This reporting stage reinforces critical thinking, strengthens oral and visual communication skills, and provides students with an authentic experience of disseminating scientific discoveries in a professional format.

## 3. Data collection

To evaluate the effectiveness of the curriculum and its impact on student learning, a comprehensive assessment strategy was implemented. Data were gathered through multiple quantitative and qualitative instruments to capture a holistic view of the student experience.

### 3.1. Anonymous Post-Course Survey

Following the completion of the module, students were invited to complete an anonymous post-course survey (**Note S4**). This survey was designed to protect student privacy while gathering honest feedback on the curriculum’s structure and delivery.

The questionnaire was divided into several key sections:

- **Background Information**: Collected data on student majors and academic levels (e.g., Freshman to Senior, or other) to understand the diversity of the cohort (**Note S4 Q1**-**3**).
- **Self-Assessment of Learning**: Measured perceived gains in conceptual understanding, research skills, technical proficiency, and bioinformatics literacy (**Note S4 Q4**-**17**).
- **Knowledge Check**: Used multiple-choice questions to objectively assess student mastery of core concepts such as the purpose of 16S rRNA sequencing, the function of PCR, and the definition of alpha diversity (**Note S4 Q18**-**22**).
- **Student Experience**: Identified the most engaging and challenging aspects of the project, as well as overall satisfaction with the learning experience (**Note S4 Q23-27**).

### 3.2. Likert-Scale Assessments

The survey utilized Likert-scale questions (typically ranging from 1 = Strongly Disagree to 5 = Strongly Agree, or Not Confident to Extremely Confident) to quantify student perceptions in the following areas:

- **Engagement**: Assessing which parts of the workflow, such as sample collection or lab work, were most stimulating for the students.
- **Confidence**: Measuring self-efficacy in designing hypothesis-driven research, processing environmental samples, and using bioinformatics tools like QIIME 2.
- **Understanding**: Evaluating the clarity of technical concepts, including the 16S rRNA gene’s role, sequencing technologies (e.g., Illumina MiSeq), and the interpretation of diversity metrics.

### 3.3. Open-Ended Reflections

To capture nuanced qualitative data, the survey included open-ended reflection questions. These prompts allowed students to describe their unique research questions and discoveries in their own words. Students also reflected on the specific skills they gained, how the experience shifted their understanding of microbiome science, and provided suggestions for future curriculum improvements.

### 3.4. Performance Artifacts

In addition to self-reported survey data, student learning was evaluated through optional performance artifacts generated during the course. These artifacts provided direct evidence of student proficiency and included:

- Lab Reports: Documenting the technical execution of the molecular workflow and PCR results.
- Scientific Posters: Summarizing the entire research cycle, from hypothesis to taxonomic classification and data visualization.

## 4. Results

Survey and assessment data were analyzed using a descriptive mixed-methods approach aligned with the four primary educational outcomes of the curriculum: (1) microbiome content knowledge, (2) research and laboratory skills, (3) computational and data-analysis literacy, and (4) scientific communication and student engagement.

### 4.1. Student Prior Experience

The curriculum was implemented with a cohort of 4 students majoring in Biochemistry (3) and Biological Science (1) (**Fig. 3A**). The majority of participants were Seniors (75%) and freshmen (25%) (**Fig. 3B**). Prior to the module, students reported varying levels of experience with molecular techniques: while all had prior exposure to PCR, 50% had experience in Genetic sequencing, and only 25% had previously worked with genetic bioinformatics tools (**Table S2**). This indicates that the module successfully introduced advanced computational and sequencing workflows to a population that was otherwise inexperienced in these areas.

**Figure 3.**
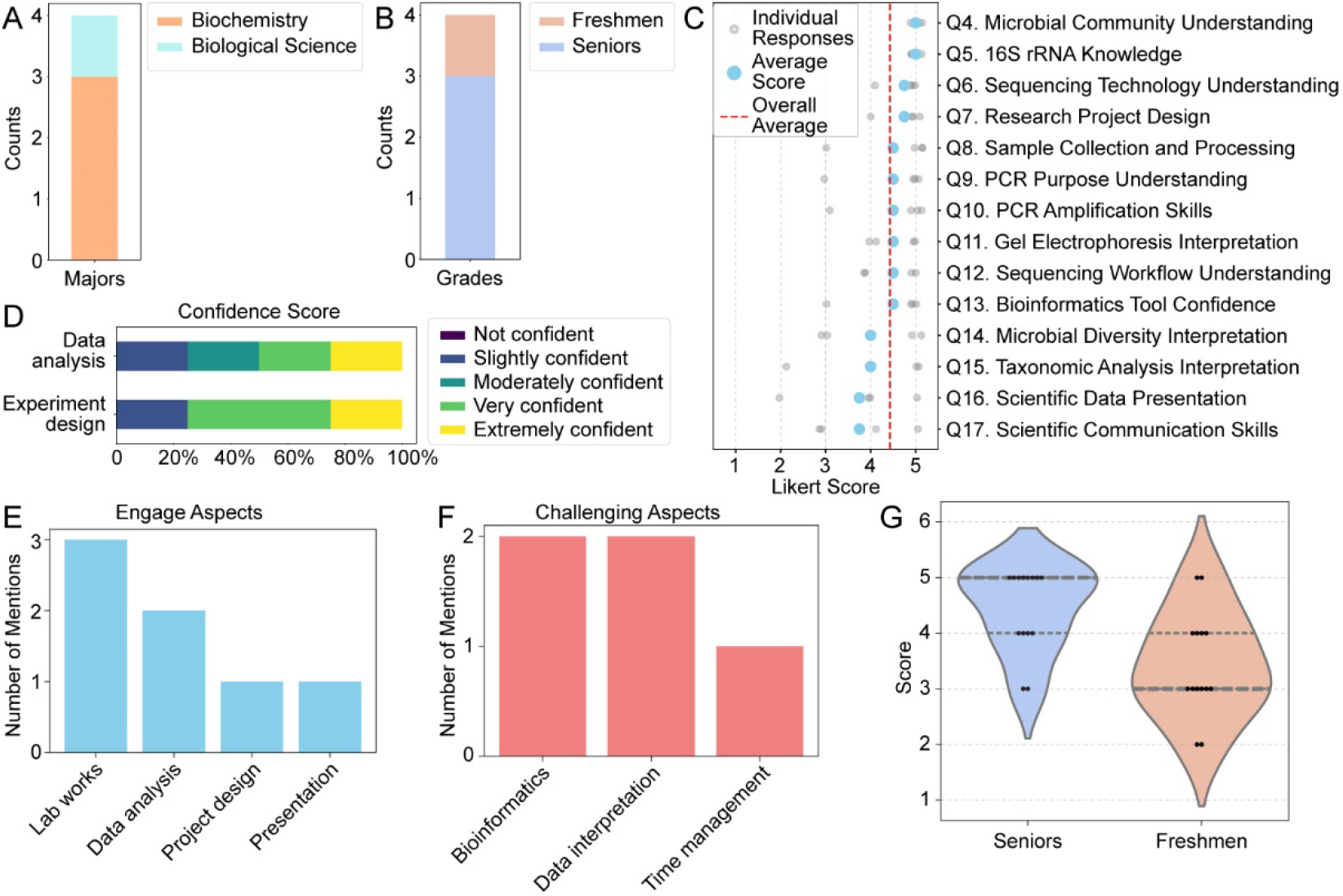
Student demographics, self-assessment, and learning outcomes. A) Student Major Distribution: Stacked bar chart showing the academic background of the participant cohort (n = 4). B) Academic Standing: Composition of the cohort by year, highlighting the participation of both Seniors and Freshmen. C) Learning Outcome Self-Assessment: Likert scale responses (1 = Strongly Disagree, 5 = Strongly Agree) across 14 categories (Q4–Q17). Small grey circles represent individual student responses; large blue circles represent the mean score for each question. The vertical red dashed line indicates the overall mean across all survey categories. D) Confidence in Core Research Pillars: Proportional confidence levels reported for Data Analysis versus Experimental Design. E) Student Engagement: Frequency of mentions for the most engaging aspects of the curriculum, with laboratory work and data analysis identified as the highest-interest areas. F) Curricular Challenges: Identification of the most difficult aspects of the module; Bioinformatics and Data Interpretation were cited as the primary hurdles for students. G) Score Distribution by Academic Level: Violin plot comparing overall performance/assessment scores between Seniors and Freshmen.

#### Student-Led Research Case Studies

A core feature of this curriculum is the transition from instructor-led protocols to student-driven inquiry. To validate the low-cost 16S rRNA workflow, students designed and executed independent investigations targeting unique local environments. Two representative projects illustrate the ability of the protocol to capture complex microbial shifts:

- *The Microbiology of Celebration (Westcott Fountain at FSU)*: One group investigated how a university tradition—throwing students into the Westcott Fountain to celebrate their 21st birthdays—affected the fountain’s water microbiome (**Poster S1**). Using a centrifugation-based cell collection and direct PCR method28, students compared microbial communities during baseline and high-activity periods. A total of 18 water samples were collected, with the estimated project cost remaining below USD 230. The study revealed pronounced microbial shifts associated with student activity and rain events. Differential abundance analysis using ANCOM-BC^[41]^identified increased levels of human-associated taxa, including *Escherichia-Shigella* and *Brevibacterium*, during peak activity periods. This project demonstrated the sensitivity and effectiveness of the low-cost workflow for detecting anthropogenic impacts on aquatic microbial community structure.
- *Impact of Disinfectants on Gym Equipment*: Another student characterized the surface microbiomes of gym equipment (dumbbells, benches, and yoga mats) and evaluated the efficacy of branded versus non-branded disinfectant wipes (**Poster S2**). Using surface swabs processed through the surfactant-assisted direct PCR protocol, students collected a total of 36 surface samples, with the estimated project cost remaining below USD 460. The analysis revealed that bacterial communities on gym equipment were dominated by skin-associated genera, including *Cutibacterium, Staphylococcus*, and *Acinetobacter*, with relatively consistent microbial compositions across different surface types. Although both disinfectant wipe types reduced microbial abundance, certain clinically relevant genera, particularly Staphylococcus and Acinetobacter, persisted following disinfection. These findings provided students with a practical demonstration of the relationship between microbial ecology, hygiene practices, and public health.

### 4.2. Validation of the Low-Cost Protocol

These student-generated results were not only educationally transformative but also scientifically rigorous. Despite the omission of expensive commercial DNA extraction kits, the surfactant-assisted direct PCR approach consistently yielded high-quality amplicons (∼400 bp) for both water and surface samples. This was verified through agarose gel electrophoresis (**Fig. S1**), where students observed clear, distinct bands ready for barcoding.

The successful generation of high-resolution data—including log relative abundance heatmaps and volcano plots (**Posters S1** and **S2**)—confirmed that the simplified laboratory workflow provides sufficient template quality for high-throughput sequencing on the Illumina MiSeq platform. By producing “conference-ready” scientific posters, students demonstrated mastery of the entire research cycle, from sampling logistics to sophisticated bioinformatic analysis in QIIME 2^[25],[26]^.

### 4.3. Learning Outcomes and Self-Assessment

Post-course surveys revealed high levels of self-reported learning across all core objectives. Using a 5-point Likert scale (1 = Strongly Disagree, 5 = Strongly Agree), students expressed high confidence in their conceptual and technical understanding:

Conceptual Understanding (**Note S4 Q4 to 6**): Students strongly agreed that they understood the role of microbial communities (mean score: 4.75) and the function of the 16S rRNA gene (mean score: 5). Students reported a moderate level of understanding of sequencing techniques (mean score = 3.75) (**Fig. 3C**).

Research and technical skills outcomes (**Note S4 Q7–Q13**) also demonstrated strong gains in student confidence across multiple competencies. Students reported the highest confidence in understanding the role of the microbiome (mean score = 4.75), performing sample collection procedures (mean score = 4.75), and understanding the purpose of PCR (mean score = 5.0). Students also expressed confidence in designing hypothesis-driven research projects (mean score = 4.5), following the sequencing workflow from sample preparation to data generation (mean score = 4.5), analyzing and interpreting taxonomic classification results (mean score = 4.5), effectively presenting scientific data (mean score = 4.5), and communicating scientific findings (mean score = 4.5) (**Fig. 3C**).

Students also reported positive levels of confidence in broader scientific research skills. When asked about experimental design, students indicated confidence levels ranging from slightly confident to extremely confident, with most students reporting that they felt very or extremely confident in designing an experiment (**Fig. 3D**). Similarly, responses related to biological data analysis ranged from slightly to extremely confident, suggesting that the course helped build student confidence in interpreting and working with biological datasets. These findings indicate that participation in the course not only improved technical understanding, but also strengthened students’ self-efficacy in conducting independent scientific research.

Knowledge Check Accuracy: Objective knowledge was validated through multiple-choice questions, where 100% of students correctly identified the primary purpose of 16S rRNA sequencing and the role of PCR in the experiment (**Table S2**).

Overall, these findings indicate that the course effectively strengthened both foundational microbiome concepts and practical research skills. High confidence in PCR, sample collection, and scientific communication suggests that the hands-on and inquiry-based learning components successfully supported student engagement and skill development. The comparatively lower confidence in sequencing techniques may reflect the technical complexity of sequencing workflows and highlights an opportunity for additional instruction or bioinformatics-focused training in future iterations of the course.

### 4.4. Student Engagement and Challenges

The “Student Experience” section highlighted that Lab Work and Data analysis were the most engaging aspects of the project, cited by 3 and 2 students, respectively (**Fig. 3E**). Conversely, Bioinformatics and Data Interpretation were identified as the most challenging aspects (2 each), suggesting that while these areas provided significant learning opportunities, they also required the most instructional support (**Fig. 3F**). Despite these challenges, the module received an “Excellent” rating from 75% of participants, and 100% of students stated they would recommend the module to others (**Table S2**).

### 4.5. Interest in Future Research

All four students expressed interest in pursuing further research in microbiology or bioinformatics after completing the project (**Table S2**). Responses highlighted interests in bacterial research, antibiotic resistance, infection detection, computational bioinformatics methods, and healthcare-related applications such as dentistry. All students also reported that the project increased their enthusiasm for research and helped connect microbiology and bioinformatics to future career goals.

### 4.6. Influence of Academic Level on Student Self-Assessment

Differences in self-reported confidence were observed between students at different academic levels. The freshman participant generally reported lower confidence scores in technical and analytical skills compared to senior students, particularly in areas related to bioinformatics, data interpretation, and independent research design (**Fig. 3G**). In contrast, senior students consistently reported higher levels of confidence across most learning objectives (**Fig. 3G**). These findings suggest that prior academic experience and exposure to advanced laboratory coursework may influence student self-evaluation and comfort with computational microbiology workflows. Nevertheless, the freshman participant still demonstrated successful engagement with the curriculum and completion of the research workflow, indicating that the module remains accessible to students at earlier stages of undergraduate training. Based on these findings, we suggest that this training module may be most effective for upper-level undergraduate students who have prior laboratory and biological science coursework experience.

## 5. Discussion

The implementation of this student-driven microbiome curriculum demonstrates that high-level genomic research can be made accessible, cost-effective, and educationally impactful for undergraduate biology and chemistry students. By shifting from a “black-box” demonstration to an authentic inquiry-based model, this module addresses critical gaps in modern life sciences education—specifically the lack of hands-on sequencing experience for students in resource-limited settings.

### 5.1. Technical Efficacy of the Low-Cost Workflow

The primary barriers to implementing 16S rRNA sequencing in the undergraduate classroom are typically the prohibitive cost of reagents and the logistical complexity of DNA extraction. Our methodology—utilizing centrifugation-based cell collection and surfactant-assisted direct PCR^[28]^—successfully bypassed these hurdles.

The student research projects (**Section 4.1**) serve as technical validation of this streamlined approach. For instance, the Westcott Fountain study demonstrated that microbial cells could be concentrated from water samples directly from the fountain without expensive vacuum filtration systems. Similarly, the Gym Equipment study showed that surfactant-assisted lysis during PCR was sufficient to amplify the DNA of the microbial cells from surface swabs. This simplified workflow reduced the per-sample cost to approximately USD 13 (**Table S1**). In comparison, conventional DNA extraction–based methods can increase costs by approximately 50%, raising the per-sample cost to nearly USD 20 (**Table S1**). In addition to lowering costs, the direct PCR workflow substantially reduced bench time and hands-on sample processing^[28]^. This allowed students to devote more attention to experimental design, hypothesis development, and data interpretation rather than repetitive laboratory procedures such as DNA extraction and extensive pipetting.

### 5.2. Impact of Student Agency and the CURE Framework

A significant finding of this study was the high level of student engagement associated with sample collection (83.3%) and project design. Unlike traditional “cookbook” laboratory exercises where outcomes are predetermined, this module provided students with the autonomy to select their own “microbial frontiers.” This agency ranged from investigating local university traditions to assessing public health hygiene.

This sense of ownership was reflected in the qualitative survey feedback, with students noting that the experience transitioned their focus from “memorizing biological facts” to “understanding the rigorous methods” of modern research. This shift is a hallmark of successful Course-based Undergraduate Research Experiences (CUREs). By contributing original data to the field of microbial ecology, students moved beyond the role of passive learners and began to develop professional scientific identities.

### 5.3. Navigating the Bioinformatics Learning Curve

While technical confidence in wet-lab skills was high, students identified bioinformatics and data interpretation as the most challenging components of the curriculum (66.7%). This difficulty was anticipated, as most participants had no prior exposure to command-line interfaces or the QIIME 2 pipeline. However, the assessment data suggests that this challenge was a catalyst for significant intellectual growth.

By grappling with Amplicon Sequence Variants (ASVs), diversity metrics, and taxonomic classification, students gained a realistic understanding of the modern biological research cycle. The fact that 100% of students recommended the module despite its perceived difficulty suggests that they found the “struggle” with authentic data to be rewarding rather than discouraging. To optimize future iterations, we suggest incorporating structured bioinformatics “dry-runs” using provided datasets to build foundational computational literacy before students analyze their independent samples.

### 5.4. Scalability and Pedagogical Alignment

The modular design of this curriculum facilitates flexible implementation across various institutional contexts. While this pilot cohort consisted of a small Directed Individual Study (DIS) group, the elimination of commercial extraction kits and the use of batched sequencing make the workflow highly scalable for larger laboratory sections of 30 or more students. Furthermore, the curriculum’s alignment with American Society for Microbiology (ASM) curricular guidelines ensures that it meets national standards for scientific thinking, quantitative analysis, and communication. This makes the module a viable, portable template for institutions seeking to integrate next-generation sequencing into their curricula without extensive infrastructure.

## 6. Conclusion

This curriculum provides a blueprint for democratizing access to next-generation sequencing in undergraduate education. By reducing the cost and technical barriers associated with traditional workflows, we enable students to act as primary investigators within their own local environments. The transition from the “Microbiology of Celebration” in a campus fountain to the “Clinical Relevance of Gym Disinfectants” illustrates that when students are equipped with the tools of modern genomics, they can transform everyday observations into sophisticated scientific inquiries. This module not only builds technical proficiency but also fosters the curiosity and critical thinking necessary for the next generation of life scientists.

## Supporting information

Poster S2

Poster S1

## Acknowledgments

This work was supported by the Florida State University startup fund (to X.L.), the Florida State University First-Year Assistant Professor Award (to X.L.), and the Innovations for Gram-Negative Antibiotic Discovery program funded by the Novo Nordisk Foundation (NNF25SA0112407 to X.L.). The authors gratefully acknowledge Drs. Amber Brown and Steven Miller at FSU for assistance with Illumina MiSeq sequencing.

## Declaration

### Consent for Publication

No individual person’s data is included in the manuscript.

## Notes

### Competing Interest Statement

The authors have declared no competing interest.

### Summary of Updates

We updated one reference, removed a duplicated session, and added the two SI posters.

