## Supplementary material for "Student-Driven Microbiome Exploration: A Low-Cost 16S rRNA Sequencing Curriculum for Undergraduate Biology Education": Supplimentary Information

**This PDF file includes:**

Figure S1: Example TapeStation trace of the 16S rRNA library

Note S1: Microbiome Research Project Proposal Template

Note S2: Laboratory Workflow for Student driven microbiome exploration: A hands-on, budget-friendly 16S sequencing curriculum.

Note S3: Bioinformatic analysis steps

Note S4: Example Master Script for Qiime 2 analysis

Note S5: Post-Course Survey Questionnaire

Table S1: Cost calculation

Table S2: Student survey and assessment data

Table S3: Primer sequences

**Other supporting materials for this manuscript include the following:**

Poster S1: The Microbiology of Celebration: How the Wescott Fountain 21st Birthday Tradition Shapes its Water Microbiome

Poster S2: Evaluating Disinfectant Wipe Impact on Gym Equipment's Surface Microbiome Using 16S rRNA Sequencing

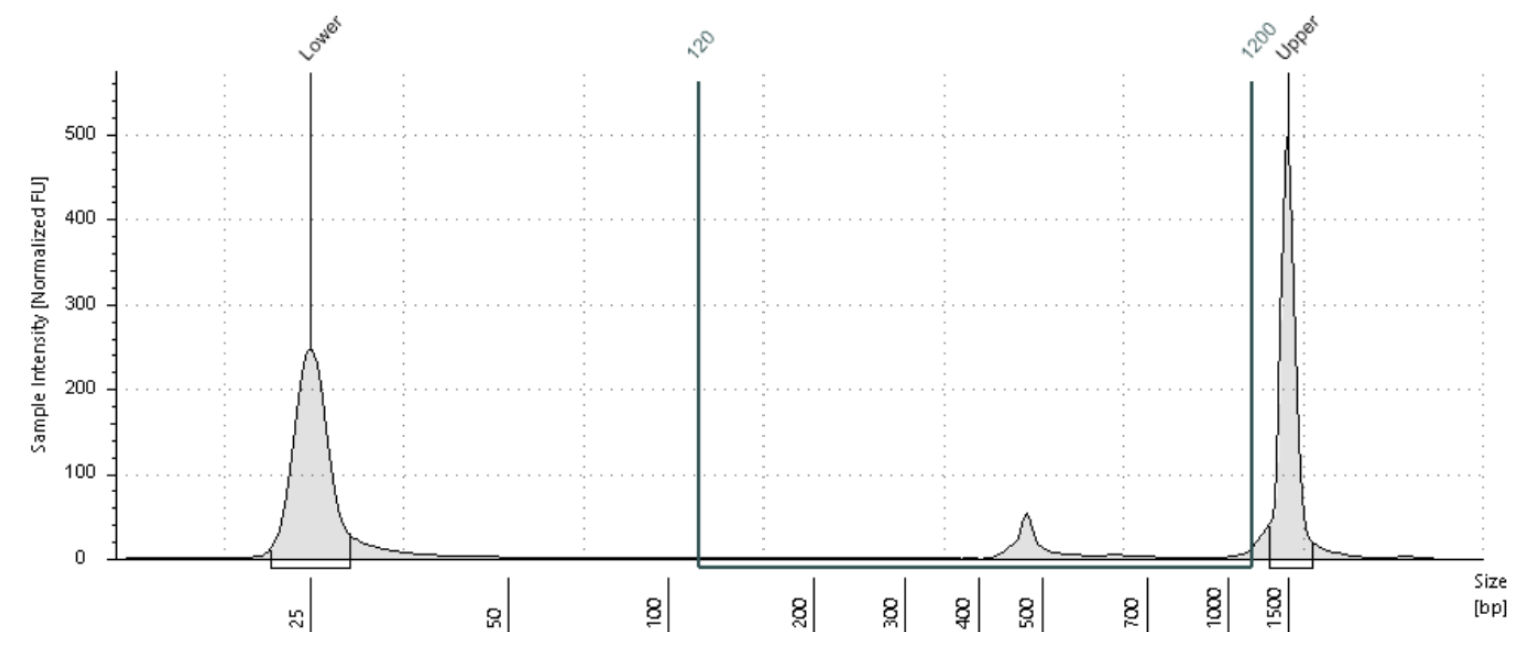

**Figure S1**: A typical TapeStation trace of the 16S rRNA library.

**Note S1: Microbiome Research Project Proposal Template**

**Project Title:**
*Provide a concise and descriptive title for your study.*

**1. Team Members and Roles**

- Student name(s):
- Roles and responsibilities of each member:

**2. Background and Rationale** (1 paragraph)

- Briefly describe the environment or surface to be studied.
- Summarize relevant concepts about microbial communities in this environment.
- Explain the significance of the study and why the question is biologically or environmentally relevant.

**3. Research Question and Hypothesis**

- **Research Question:**
  *State the main question your project aims to address.*
- **Hypothesis:**
  *Provide a clear, testable hypothesis and expected outcome.*

**4. Experimental Design** (Answer the questions)

**4.1 Sampling Plan**

- Sampling location(s):
- Sample type (e.g., water, surface swab):
- Number of samples and replicates:
- Sampling schedule (date/time or frequency):

**4.2 Controls**

- Negative control(s):
- Positive control(s) (if applicable):
- Rationale for chosen controls:

**4.3 Variables**

- Independent variable(s):
- Dependent variable(s):
- Controlled variables:

**5. Methods Overview** (DO NOT CHANGE THIS SECTION)

- Sample collection procedure (brief description).
- Microbial cell collection (centrifugation and washing).
- Direct PCR amplification of the 16S rRNA gene (V3–V4 region).
- Library preparation and sequencing approach.
- Bioinformatics analysis plan (e.g., QIIME 2, diversity metrics, taxonomic profiling).

**6. Data Analysis Plan**

- Describe how sequencing data will be processed and analyzed.
- Planned comparisons (e.g., between locations, surfaces, or time points).
- Types of outputs to be generated (e.g., taxonomic bar plots, α- and β-diversity, ordination plots).

**7. Feasibility and Resources**

- Sample accessibility:
- Estimated number of samples:

**8. Biosafety and Ethical Considerations**

- Potential biosafety risks associated with sampling:
- Required permissions or approvals for sampling locations:
- Planned safety procedures and contamination-control practices:

**9. Expected Outcomes and Potential Limitations**

- Anticipated results based on the hypothesis:
- Possible challenges or limitations (e.g., low biomass, contamination, sequencing depth).
- Strategies to mitigate these limitations.

**10. Project Timeline**

| **Step** | **Planned Date** | **Notes** |
| --- | --- | --- |
| Project design finalized |  |  |
| Sample collection |  |  |
| PCR and library preparation |  |  |
| Sequencing submission |  |  |
| Bioinformatics analysis |  |  |
| Poster preparation |  |  |

**This is the end of Note S1**

**Note S2: Laboratory Workflow for:** Student driven microbiome exploration: A hands-on, budget-friendly 16S sequencing curriculum.

**Objectives:**

- Explain the role of microbial communities in natural and engineered environments and their relevance to human and ecosystem health.
- Design and conduct an independent microbiome study, including hypothesis formulation, sampling strategy, and experimental planning.
- Apply molecular biology techniques such as DNA extraction and 16S rRNA gene amplification to characterize microbial communities.
- Analyze and interpret sequencing data using open-source bioinformatics pipelines (e.g., QIIME 2) to evaluate microbial diversity and taxonomy.
- Communicate scientific findings effectively through written reports or poster presentations.

**Project design:**

1. Attend an introductory lecture on microbiome research and 16S rRNA gene sequencing, covering microbial diversity, taxonomic profiling, and hypothesis-driven investigation.
2. Identify a research topic related to microbial communities in everyday environments (e.g., swimming pools, rivers, or shared surfaces such as buses, gym equipment, and library computer keyboards).
3. Formulate a clear, testable hypothesis based on the selected research question (e.g., the river microbiome is influenced by seasonal changes).
4. Develop an experimental plan that includes the sampling strategy, appropriate controls, and analytical approach; submit the written proposal to the professor and graduate teaching assistants. The template of the written proposal is provided (see **Supplementary Information**).
5. Evaluate feasibility, resource requirements, and alignment with the low-cost workflow.
6. The professor and graduate teaching assistants review the proposal to assess feasibility, resource needs, biosafety risks, ethical considerations, and appropriate safety and contamination-control practices, and then provide structured feedback.
7. Revise and refine the proposal based on instructor guidance, class discussions, and iterative feedback.
8. Finalize a scientifically sound and feasible study design for implementation in subsequent laboratory sessions.

**Procedure:**

1. Cell collection from water samples

Water Samples: Water samples are collected in sterile 50 mL centrifuge tubes and immediately transported on ice to the laboratory for processing. Samples are centrifuged at 4000 × g for 10 min to pellet microbial cells. The supernatant is carefully removed without disturbing the pellet, leaving approximately 1 mL of residual liquid. This remaining volume is transferred to a 1.5 mL microcentrifuge tube and centrifuged again to further concentrate the cells. The pellet is washed three times with 1 mL of Tris buffer, retaining ~50 µL after each wash. Finally, the sample is transferred to PCR tubes, centrifuged once more, and most of the supernatant is removed, leaving ~5 µL of liquid. The pellet may not be visible at each step. Samples can be stored at −80 °C for long-term preservation.

Surface Samples: Surface samples are collected using sterile cotton swabs pre-wetted with Tris HCl buffer containing 0.1% Tween 20. The defined surface is wiped using a rotational motion to maximize cell recovery. The swab is then transferred to a 1.5 mL tube containing the same buffer, soaked, and gently squeezed several times to release bacterial cells into the solution. Samples are transported on ice to the laboratory and processed immediately. Tubes are centrifuged at 4000 × g for 10 min to pellet the cells. The supernatant is carefully removed without disturbing the pellet, leaving approximately 50 µL of liquid. The suspension is transferred to a 1.5 mL microcentrifuge tube and centrifuged again. The pellet is washed three times with 1 mL of Tris buffer, retaining ~50 µL after each wash. The final suspension is transferred to PCR tubes, centrifuged once more, and the supernatant is removed, leaving ~5 µL. The pellet may not be visible during processing. A swing-bucket centrifuge is recommended to improve pellet recovery compared to a fixed-angle rotor. Samples can be stored at −80 °C for extended periods.

2. Direct PCR for 16S rRNA amplification

The cells in the PCR tubes are mixed with the PCR reagent as follow:

| emulsion PCR | stock | final | Volume (uL) |
| --- | --- | --- | --- |
| Q5 Master mix | 2 | 1 | 25 |
| Nx_515F (uM) | 4 | 0.4 | 5 |
| Nx_806R (uM) | 4 | 0.4 | 5 |
| IGEPAL | 1% | 0.10% | 5 |
| Cells from last step | |  | 5 |
| H2O |  |  | 5 |
| total |  |  | 50 |

The tubes are thermocycled with the following program: (95 °C 30 s; **35** cycles of 95 °C 5 s, 55 °C 30 s, 72 °C 1.5 min; 72 °C 2 min; 12 °C hold).

3. Amplicon purification by size selection

AMPure XP beads (or similar products) are added to PCR tubes in a ratio of 0.7 µL AMPure XP beads per 1 µL of PCR product (for the 50 µL PCR, 35 µL AMPure XP bead are added). All mixing steps were completed by gentle vertexing or flicking. Upon addition of the AMPure XP beads, the solution is mixed, and incubated for 5 min at room temperature. Two ethanol washes following magnetic separation were performed with 500 μL of 80% EtOH, and then the beads were air-dried for 5-10 min. The DNA are then eluted with 16 µL 10 mM Tris-HCl pH 7.5 buffer or Nuclease free water. Amplicon QC are performed using Qubit (for concentration) and TapeStation (for size distribution). The amplicons can be stored at -20 for long term.

4. Index PCR for sequencing library preparation

The amplicon from previous step are diluted to 1 ng/μL, and mixed with the PCR mix as follow:

If multiple samples will be mixed for sequencing on the same lane, they should use different indices.

| Index PCR | stock | final | volume |
| --- | --- | --- | --- |
| Q5 Master mix | 2 | 1 | 25 |
| Nx7xx (uM) | 4 | 0.4 | 5 |
| Nx5xx (uM) | 4 | 0.4 | 5 |
| DNA | 1 |  | 5 |
| H2O |  |  | 10 |
| total |  |  | 50 |

The tubes are thermocycled with the following program: (95 °C 30 s; **8** cycles of 95 °C 5 s, 55 °C 30 s, 72 °C 1.5 min; 72 °C 2 min; 12 °C hold).

5. Library purification

The library is purified using similar process as step 3 (Amplicon purification by size selection) and eluted into 12 μL. After QC, the library is mixed and sequenced according to Illumina Miseq protocol.

**This is the end of Note S2**

**Note S3:** Bioinformatic analysis steps

To ensure the pipeline runs smoothly, your input files must follow the strict formatting rules required by QIIME 2. Below is a guide on how to prepare your **Manifest** and **Metadata** files.

**1. Preparing the Manifest File**

The manifest file tells QIIME 2 where your raw Fastq files are located and which sample ID belongs to which file. It is a tab-separated file that must use specific headers depending on your data type. For Single-End, use two columns: sample-id and absolute-filepath. For Paired-End, use three: sample-id, forward-absolute-filepath, and reverse-absolute-filepath. Always ensure you use absolute paths (starting with /) to prevent "file not found" errors.

**Example Manifest:**

**For pair-end reads:**

| sample-id | forward-absolute-filepath | reverse-absolute-filepath |
| --- | --- | --- |
| sample-1 | $PWD/some/filepath/sample0_R1.fastq.gz | $PWD/some/filepath/sample1_R2.fastq.gz |
| sample-2 | $PWD/some/filepath/sample2_R1.fastq.gz | $PWD/some/filepath/sample2_R2.fastq.gz |
| sample-3 | $PWD/some/filepath/sample3_R1.fastq.gz | $PWD/some/filepath/sample3_R2.fastq.gz |
| sample-4 | $PWD/some/filepath/sample4_R1.fastq.gz | $PWD/some/filepath/sample4_R2.fastq.gz |

**For single-end reads**

| sample-id | absolute-filepath |
| --- | --- |
| sample-1 | $PWD/some/filepath/sample1_R1.fastq |
| sample-2 | $PWD/some/filepath/sample2_R1.fastq |

**Pro Tip:** You can create this easily in Excel and "Save As" **Tab Delimited Text (.txt)**, then rename the extension to .tsv.

**2. Preparing the Metadata File**

The metadata file contains the experimental variables for your samples, such as treatment groups or dates. The first column must be named sample-id and must exactly match the IDs in your manifest. When naming your other columns, avoid spaces and special characters; instead, use underscores (e.g., *treatment_group*) to ensure compatibility with R-based plugins like ANCOM-BC.

**Rules:**

- **Format:** Tab-Separated Value (.tsv).
- **Required Header:** The first column **must** be sample-id.
- **Case Sensitivity:** QIIME 2 is case-sensitive. "Before" and "before" will be treated as different groups.
- **ANCOM-BC Note:** Avoid using special characters (like #, @, +, or spaces) in your column names or cell values. Use underscores (_) instead (e.g., wipe_brand instead of Wipe Brand).

**Example Metadata:**

| sample-id | wipe_brand | equipment_type | time_point |
| --- | --- | --- | --- |
| HB_GE_1 | branded | treadmill | before |
| HB_GE_2 | branded | treadmill | after |
| HB_GE_3 | non-branded | dumbbell | before |

**3. How to Use with the Master Script**

Once your manifest and metadata files are prepared and uploaded, follow these steps to configure and launch the pipeline:

1. **Upload Files**: Place your manifest.tsv and metadata.tsv onto your server. It is recommended to keep them in a dedicated folder, such as 01_manifest_metadata, to keep your project workspace organized.
2. **Edit the Master Script**: Open your master_pipeline.sh file with a text editor (like nano or vi) and navigate to the **User Configuration** section at the top.
3. **Set File Paths**: Update the following variables with the **absolute paths** to your files:

MANIFEST="/path/to/your/01_manifest_metadata/manifest.tsv"

METADATA="/path/to/your/01_manifest_metadata/metadata.tsv"

4. **Configure Parameters**:
   - **SAMPLING_DEPTH**: Enter your desired rarefaction depth. This is typically based on the minimum frequency of reads per sample (your previous gym microbiome analysis used **18,058**).
   - **EXCLUDE_SAMPLES**: If you identified low-quality samples that need to be dropped (such as HB_GE_22 or HB_GE_4), list them here using the pipe (|) symbol as a separator.
5. **Verify Metadata**: Before committing to a full run, it is a best practice to verify that QIIME 2 can read your metadata without errors. Run this command to generate a summary visualization:

qiime metadata tabulate --m-input-file your_metadata.tsv --o-visualization check.qzv

View the resulting .qzv file at [view.qiime2.org](https://view.qiime2.org) to ensure all columns and sample IDs are correctly formatted.

6. **Run the Pipeline**: Once verified, turn on the qiime2 environment and execute the script from your terminal:

conda activate qiime2-amplicon-2024.2

./master_pipeline.sh

**Note S4: Example Master Script for Qiime 2 analysis**

Copy the following text in green into a new txt file, edit the values, and save it as ‘master_pipeline.sh’

#!/bin/bash

\# ==============================================================================

\# Generalized QIIME 2 Automated Pipeline

\# ==============================================================================

set -e # Exit immediately if a command exits with a non-zero status

set -u # Treat unset variables as an error

\# ------------------------------------------------------------------------------

\# 1. USER CONFIGURATION (Edit these values)

\# ------------------------------------------------------------------------------

PROJECT_NAME="gym_microbiome_reanalysis"

BASE_DIR="/data10/hassan/${PROJECT_NAME}"

\# Input File Paths

MANIFEST="/data10/hassan/gym_microbiome_reanalysis/01_manifest_metadata/manifest.tsv"

METADATA="/data10/hassan/gym_microbiome_reanalysis/01_manifest_metadata/metadata.tsv"

CLASSIFIER="/data10/hassan/LC_project/data/silva-138-99-nb-classifier.qza"

\# Pipeline Parameters

SAMPLING_DEPTH=18058

TRUNC_LEN=131

THREADS=0 # '0' uses all available cores

\# Sample Filtering (IDs to remove, separated by | for regex)

EXCLUDE_SAMPLES="HB_GE_22|HB_GE_4"

\# ------------------------------------------------------------------------------

\# 2. SETUP DIRECTORIES & LOGGING

\# ------------------------------------------------------------------------------

echo "Creating project structure at ${BASE_DIR}..."

mkdir -p ${BASE_DIR}/{02_import,03_cutadapt/logs,03_quality,04_dada2,05_taxonomy,06_phylogeny,07_diversity/logs,08_ancombc/logs}

\# Define derived metadata paths

METADATA_FILTERED="${BASE_DIR}/01_manifest_metadata/metadata_filtered.tsv"

\# ------------------------------------------------------------------------------

\# 3. IMPORT & QUALITY CONTROL

\# ------------------------------------------------------------------------------

echo "Step 02: Importing data..."

qiime tools import \

--type 'SampleData[SequencesWithQuality]' \

--input-path "$MANIFEST" \

--output-path "${BASE_DIR}/02_import/demux.qza" \

--input-format SingleEndFastqManifestPhred33V2

echo "Step 03: Trimming primers and checking quality..."

qiime cutadapt trim-single \

--i-demultiplexed-sequences "${BASE_DIR}/02_import/demux.qza" \

--p-front GTGYCAGCMGCCGCGGTAA \

--p-discard-untrimmed \

--p-minimum-length 100 \

--o-trimmed-sequences "${BASE_DIR}/03_cutadapt/demux_trimmed.qza" \

--output-dir "${BASE_DIR}/03_cutadapt/logs" \

--verbose

qiime demux summarize \

--i-data "${BASE_DIR}/02_import/demux.qza" \

--o-visualization "${BASE_DIR}/03_quality/demux_summary.qzv"

\# ------------------------------------------------------------------------------

\# 4. DENOISING (DADA2)

\# ------------------------------------------------------------------------------

echo "Step 04: Running DADA2 denoising..."

qiime dada2 denoise-single \

--i-demultiplexed-seqs "${BASE_DIR}/03_cutadapt/demux_trimmed.qza" \

--p-trim-left 0 \

--p-trunc-len $TRUNC_LEN \

--p-n-threads $THREADS \

--o-table "${BASE_DIR}/04_dada2/feature_table.qza" \

--o-representative-sequences "${BASE_DIR}/04_dada2/rep_seqs.qza" \

--o-denoising-stats "${BASE_DIR}/04_dada2/denoising_stats.qza" \

--o-base-transition-stats "${BASE_DIR}/04_dada2/base_transition_stats.qza" \

--verbose 2>&1 | tee "${BASE_DIR}/04_dada2/dada2_log.txt"

\# ------------------------------------------------------------------------------

\# 5. TAXONOMY & FILTERING

\# ------------------------------------------------------------------------------

echo "Step 05: Taxonomic classification and filtering..."

qiime feature-classifier classify-sklearn \

--i-classifier "$CLASSIFIER" \

--i-reads "${BASE_DIR}/04_dada2/rep_seqs.qza" \

--o-classification "${BASE_DIR}/05_taxonomy/taxonomy.qza" \

--p-n-jobs $THREADS

\# Filter Table: Remove non-bacterial/contaminant sequences

qiime taxa filter-table \

--i-table "${BASE_DIR}/04_dada2/feature_table.qza" \

--i-taxonomy "${BASE_DIR}/05_taxonomy/taxonomy.qza" \

--p-exclude mitochondria,chloroplast,Eukaryota,Archaea,Unassigned \

--p-include p__ \

--o-filtered-table "${BASE_DIR}/05_taxonomy/feature_table_filtered.qza"

\# Filter Samples: Remove specified IDs and update metadata

qiime feature-table filter-samples \

--i-table "${BASE_DIR}/05_taxonomy/feature_table_filtered.qza" \

--m-metadata-file "$METADATA" \

--p-exclude-ids \

--p-where "[sample-id] IN ('${EXCLUDE_SAMPLES//|/\',\'}')" \

--o-filtered-table "${BASE_DIR}/05_taxonomy/feature_table_final.qza"

grep -vE "$EXCLUDE_SAMPLES" "$METADATA" > "$METADATA_FILTERED"

\# ------------------------------------------------------------------------------

\# 6. PHYLOGENY & DIVERSITY

\# ------------------------------------------------------------------------------

echo "Step 06: Generating phylogenetic tree..."

qiime phylogeny align-to-tree-mafft-fasttree \

--i-sequences "${BASE_DIR}/04_dada2/rep_seqs.qza" \

--o-alignment "${BASE_DIR}/06_phylogeny/aligned.qza" \

--o-masked-alignment "${BASE_DIR}/06_phylogeny/masked.qza" \

--o-tree "${BASE_DIR}/06_phylogeny/unrooted_tree.qza" \

--o-rooted-tree "${BASE_DIR}/06_phylogeny/rooted_tree.qza" \

--p-n-threads $THREADS

echo "Step 07: Diversity Analysis..."

qiime diversity core-metrics-phylogenetic \

--i-phylogeny "${BASE_DIR}/06_phylogeny/rooted_tree.qza" \

--i-table "${BASE_DIR}/05_taxonomy/feature_table_final.qza" \

--p-sampling-depth $SAMPLING_DEPTH \

--m-metadata-file "$METADATA_FILTERED" \

--output-dir "${BASE_DIR}/07_diversity/core_metrics"

\# ------------------------------------------------------------------------------

\# 7. DIFFERENTIAL ABUNDANCE (ANCOM-BC)

\# ------------------------------------------------------------------------------

echo "Step 08: Running ANCOM-BC..."

qiime taxa collapse \

--i-table "${BASE_DIR}/05_taxonomy/feature_table_final.qza" \

--i-taxonomy "${BASE_DIR}/05_taxonomy/taxonomy.qza" \

--p-level 6 \

--o-collapsed-table "${BASE_DIR}/08_ancombc/table_genus.qza"

qiime composition ancombc \

--i-table "${BASE_DIR}/08_ancombc/table_genus.qza" \

--m-metadata-file "$METADATA_FILTERED" \

--p-formula "wipe_brand+equipment_type+time_point" \

--o-differentials "${BASE_DIR}/08_ancombc/ancombc_results.qza"

echo "=========================================="

echo "Pipeline Complete for ${PROJECT_NAME}"

echo "=========================================="

**This is the end of Note S4**

**Note S5: Post-Course Survey Questionnaire**

**Section 1: Background Information**

1. What is your major?
2. What is your academic level?
   - Freshman
   - Sophomore
   - Junior
   - Senior
   - Other
3. Prior to this course, had you worked with (Check all that apply):
   - PCR
   - Genetic sequencing
   - Bioinformatics tools (e.g., QIIME2)

**Section 2: Self-Assessment of Learning**

*All questions in this section use a scale of 1 (Strongly Disagree) to 5 (Strongly Agree).*

*Conceptual Understanding*

4. I understand the role of microbial communities in environmental and biological systems.
5. I can explain what the 16S rRNA gene is and why it is used in microbiome studies.
6. I understand how sequencing technologies (e.g., Illumina MiSeq) generate microbial data.

*Research Skills*

7. I can design a hypothesis-driven microbiome research project.
8. I feel confident collecting and processing environmental samples.
9. I understand the purpose of PCR in microbial analysis.

*Technical Skills*

10. I can perform or explain the steps of PCR amplification.
11. I understand how to interpret agarose gel electrophoresis results.
12. I can follow a sequencing workflow from sample to data output.

*Bioinformatics & Data Analysis*

13. I feel confident using bioinformatics tools such as QIIME 2.
14. I can interpret microbial diversity metrics (e.g., alpha and beta diversity).
15. I can analyze and interpret taxonomic classification results.

*Scientific Communication*

16. I can effectively present scientific data in visual formats (graphs, charts).
17. I can clearly communicate scientific findings in writing or presentations.

**Section 3: Knowledge Check (Multiple Choice)**

18. What is the primary purpose of 16S rRNA sequencing?

* To sequence entire genomes

* To identify and compare microbial communities

* To amplify proteins

* To measure RNA expression

19. What is PCR used for in this experiment?

* Protein synthesis

* DNA amplification

* Cell growth

* Sequencing directly

20. Which step comes first in a microbiome workflow?

* Data analysis

* PCR amplification

* Sample collection

* Taxonomic classification

21. What does "alpha diversity" measure?

* Differences between samples

* Diversity within a sample

* DNA concentration

* Sequencing errors

22. Which tool is commonly used for microbiome data analysis?

* Microsoft Excel

* QIIME 2

* Photoshop

* Kbase

**Section 4: Skills & Confidence**

23. How confident are you in designing an experiment?

* Not confident

* Slightly confident

* Moderately confident

* Very confident

* Extremely confident

24. How confident are you in analyzing biological data?

* Not confident

* Slightly confident

* Moderately confident

* Very confident

* Extremely confident

**Section 5: Student Experience**

25. What part of the project did you find most engaging? (Check all that apply)

* Project design

* Sample collection

* Lab work (PCR, gels)

* Data analysis

* Presentation

26. What was the most challenging aspect of the module? (Check all that apply)

* Experimental design

* Lab techniques

* Bioinformatics

* Data interpretation

* Time management

**Section 6: Open-Ended Reflection**

27. Describe your research question and what you discovered.
28. What skills did you gain from this module?
29. How has this experience changed your understanding of microbiome science?
30. What improvements would you suggest for this curriculum?
31. Would you be interested in pursuing further research in microbiology or bioinformatics? Why or why not?

**Section 7: Overall Evaluation**

32. Overall, how would you rate this learning experience?

* Poor

* Fair

* Good

* Very Good

* Excellent

33. Would you recommend this module to other students?

* Yes

* No

**This is the end of Note S5**

Table S1: Cost calculation

| **Reagents** | **vendor** | **catalog #** | **price** | **reactions** | **per sample cost** |
| --- | --- | --- | --- | --- | --- |
| Q5® High-Fidelity 2X Master Mix | New England Biolabs | M0492L | 898 | 500 | 3.592 |
| Tris buffer | Sigma | T2194-100ML | 40.1 | 200 | 0.2005 |
| Tween 20 | Sigma Aldrich | P2287100ML | 66.55 | 10000 | 0.006655 |
| 50 mL tube | VWR | 77587-932 | 98.43 | 300 | 0.3281 |
| 1.5 mL tube | VWR | 76332-064 | 36.77 | 500 | 0.07354 |
| PCR tubes | VWR | 20170-010 | 40.59 | 1000 | 0.04059 |
| Primer* | IDT DNA | See **Table S3** | 2503.2 | 15000 | 0.16688 |
| Igepal | VWR | IC0219859680 | 79.28 | 200000 | 0.000396 |
| Ampure XP | VWR | MSPP-A63881 | 2198.57 | 1200 | 1.832142 |
| EtOH | WWR | 71001-754 | 87.23 | 10000 | 0.008723 |
| Water, Molecular Biology Grade | VWR | VWRL0201-1000 | 25.83 | 1000 | 0.02583 |
| Tapestation Screen tape | Agilent Technologies | 50675582 | 344.5 | 112 | 3.075893 |
| MiSeq kit** | Illumina | MS-102-2003 | 1661 | 500 | 3.322 |
| **Estimated cost** |  |  |  |  | **12.67325** |
| If DNA extraction kit was used |  |  |  |  |  |
| Power soil DNA extraction kit | Qiagen | 12855-100 | 694.4 | 100 | 6.944 |
| **Estimated cost with DNA extraction** | |  |  |  | **19.61725** |

* Estimated based on IDT 250 nmol oligonucleotide ordering costs at approximately USD 1.49 per base for 24 forward and 16 reverse index primers, along with two 16S primers, with a guaranteed yield of 60 nmol.

** Calculated based on 12 million reads per kit, with each sample requiring at least 24K reads; therefore, each kit can be used to sequence approximately 500 samples.

Table S2: Student survey and assessment data

| 1. What is your major? | Biochemistry | Biochemistry | Biochemistry | Biological Science |
| --- | --- | --- | --- | --- |
| 2. What is your academic level? | Senior | Freshman | Senior | Senior |
| 3. Prior to this course, had you worked with: | PCR | PCR | PCR, Genetic sequencing | PCR, Genetic sequencing, Bioinformatics tools (e.g., QIIME2) |
| 4. I understand the role of microbial communities in environmental and biological systems. | 5 | 4 | 5 | 5 |
| 5. I can explain what the 16S rRNA gene is and why it is used in microbiome studies. | 5 | 5 | 5 | 5 |
| 6. I understand how sequencing technologies (e.g., Illumina MiSeq) generate microbial data. | 3 | 3 | 5 | 4 |
| 7. I can design a hypothesis-driven microbiome research project. | 5 | 3 | 5 | 5 |
| 8. I feel confident collecting and processing environmental samples. | 5 | 4 | 5 | 5 |
| 9. I understand the purpose of PCR in microbial analysis. | 5 | 5 | 5 | 5 |
| 10. I can perform or explain the steps of PCR amplification. | 4 | 4 | 5 | 5 |
| 11. I understand how to interpret agarose gel electrophoresis results. | 3 | 3 | 5 | 5 |
| 12. I can follow a sequencing workflow from sample to data output. | 4 | 4 | 5 | 5 |
| 13. I feel confident using bioinformatics tools such as QIIME 2. | 4 | 2 | 5 | 4 |
| 14. I can interpret microbial diversity metrics (e.g., alpha and beta diversity). | 4 | 2 | 5 | 5 |
| 15. I can analyze and interpret taxonomic classification results. | 5 | 3 | 5 | 5 |
| 16. I can effectively present scientific data in visual formats (graphs, charts). | 5 | 3 | 5 | 5 |
| 17. I can clearly communicate scientific findings in writing or presentations. | 5 | 3 | 5 | 5 |
| 18. What is the primary purpose of 16S rRNA sequencing? | To identify and compare microbial communities | To identify and compare microbial communities, To measure RNA expression | To identify and compare microbial communities | To identify and compare microbial communities |
| 19. What is PCR used for in this experiment? | DNA amplification | DNA amplification, Sequencing directly | DNA amplification | DNA amplification |
| 20. Which step comes first in a microbiome workflow? | Sample collection | Sample collection | Sample collection | Sample collection |
| 21. What does “alpha diversity” measure? | Diversity within a sample | Diversity within a sample | Diversity within a sample | Diversity within a sample |
| 22. Which tool is commonly used for microbiome data analysis? | QIIME 2 | QIIME 2, Kbase | QIIME 2 | QIIME 2 |
| 23. How confident are you in designing an experiment? | Very confident | Slightly confident | Extremely confident | Very confident |
| 24. How confident are you in analyzing biological data? | Slightly confident | Moderately confident | Extremely confident | Very confident |
| 25. What part of the project did you find most engaging? | Lab work (PCR, gels) | Lab work (PCR, gels), Data analysis | Project design, Data analysis | Lab work (PCR, gels), Presentation |
| 26. What was the most challenging aspect of the module? | Bioinformatics | Experimental design, Data interpretation, Time management | Experimental design, Data interpretation | Bioinformatics |
| 27. Describe your research question and what you discovered. | My research question focused on how microbial communities differ across gym equipment surfaces. I found that gym equipment harbors similar microbial populations with slight variations that can be due to different surfaces. Also, we compared the effects of two types of wipes on microbial compositions and we found that both wipes uniformly removed different types of bacteria equally. | Does student activity change the bacteria within the water fountains in student dormitories? Data inconclusive as of April 15, 2026 | Mentored undergraduate students in independent studies on the amplicon sequence analysis of 16S rRNA genes, each addressing a unique biological hypothesis. My involvement included overseeing the entire QIIME2 analysis pipeline and subsequent R plotting. All students succeeded in identifying community differences regardless of their individual sample numbers. | The Microbiology of Celebration: How the Wescott Fountain 21st Birthday Tradition Shapes its Water Microbiome. For this experiment, I discovered that student fountain throwing significantly enriches human associated taxa in the water fountain. Rain events showed to dilute the resident microbiome. Time of day had no independent effect on community composition. During active periods, 61 genera were significantly enriched including skin, gut, and oral associated taxa indicative of human activity.  Genome Analysis of a Multi-Antimicrobial Resistant Leucobacter Isolate (Q13C). For this experiment, it was discovered that 14 genes were annotated to antimicrobial drug resistance. Genotypic analysis identified resistance genes across five antibiotic classes but phenotypic disk diffusion was important towards linking these results. This isolate had confirmed resistance across four antibiotic classes when validated by both genotypic and phenotypic evidence. Bioinformatic annotation requires phenotypic confirmation and that gene presence alone is not sufficient for accurate resistance predictions. |
| 28. What skills did you gain from this module? | I gained experience in microbiome sampling, PCR, 16S rRNA sequencing workflow, data interpretation, and using bioinformatics tools such as QIIME 2. | Laboratory skills, sample collection, experimental desigm. | Improved teaching skills for computational biology for beginners by presenting complicated bioinformatics algorithms through simplified steps. Enhanced knowledge about QIIME2 workflow and R graphics by explaining every choice I made in a way that a newcomer would understand. | Data Analysis, Lab Techniques, Experimental Design, Bioinformatics, and Presentation |
| 29. How has this experience changed your understanding of microbiome science? | This experience showed me how complex microbial communities are and how sequencing can be used to study microbes in everyday environments. In our project specifically, it also demonstrated how resilient these microbial communities can be, even on frequently cleaned and heavily used gym equipment. | It has shown me how complex and multifaceted it truly is. As well, I now know how many moving parts are needed to fully analyze the environment. | Seeing several students go through the process using the same pipeline but on different data sets made me realize how important the analytical steps are to determine the outcome of the analysis. | Transitioning from a a lecture class to a lab environment shows that microbiome science is a meticulous process where small technical errors in DNA extraction or PCR can significantly alter your data. Hands-on research demonstrates how environmental factors directly influence microbial populations, turning abstract concepts like diversity into observable patterns. Analyzing real sequencing data through bioinformatics tools provides a deeper appreciation for the immense complexity of these micro communities. Ultimately, I feel like this experience shifts your focus from simply memorizing biological facts to really understanding the rigorous methods used to study the natural world. I have gained so much knowledge in this area by simply being in this lab doing hands on work while being able to analyze data that I performed in the lab. |
| 30. What improvements would you suggest for this curriculum? | Adding more step-by-step guidance for the bioinformatics analysis portion would make the curriculum easier to follow. | Give a little bit more information to the students to make their projects | A session on basic command line steps before embarking on QIIME2, spending time understanding the intermediate outputs, and spending enough time on visualizing using R. | I feel like there wasn’t any improvements needed. The structure and layout of this curriculum made it easy for us students to work at their own pace while getting the guidance they needed in areas they felt least confident on. It was a bridge between choosing an experiment that was fun while still learning the essentials behind microbial communities. |
| 30. What improvements would you suggest for this curriculum? 2 | Adding more step-by-step guidance for the bioinformatics analysis portion would make the curriculum easier to follow. | N/A | A session on basic command line steps before embarking on QIIME2, spending time understanding the intermediate outputs, and spending enough time on visualizing using R. | I feel like there wasn’t any improvements needed. The structure and layout of this curriculum made it easy for us students to work at their own pace while getting the guidance they needed in areas they felt least confident on. It was a bridge between choosing an experiment that was fun while still learning the essentials behind microbial communities. |
| 31. Would you be interested in pursuing further research in microbiology or bioinformatics? Why or why not? | Yes, because this project increased my interest in microbiology and showed me how powerful bioinformatics can be for studying complex biological systems. | Absolutely. I want to work with bacteria, either in infection detection, drug discovery, or ABR. This like of work is fascinating and I want to continue pursuing it. | Yes, especially in bioinformatics. The process of mentoring made me realize how much of a need there was for good, well-organized computational pipelines, and I am interested in developing this further. | I think microbiology is a field I would definitely pursue further research in because it is such an important topic in day to day life. I am pursuing a career in dentistry where microbes in the mouth or on the teeth are studied in tooth decay and biofilm formation. Even antibiotic resistant microbes are super important to study due to a world leading towards super mutant bacteria since microbes build resistance towards common antibiotics ingested by humans. |
| 32. Overall, how would you rate this learning experience? | Excellent | Very Good | Excellent | Excellent |
| 33. Would you recommend this module to other students? | Yes | Yes | Yes | Yes |

Table S3: Primer sequences

| nx_515F | TCGTCGGCAGCGTCAGATGTGTATAAGAGACAG GTGYCAGCMGCCGCGGTAA |
| --- | --- |
| nx_806R | GTCTCGTGGGCTCGGAGATGTGTATAAGAGACAG GGACTACNVGGGTWTCTAAT |
| index primers (nextera) N7xx | |
| 701 | CAAGCAGAAGACGGCATACGAGAT TCGCCTTA GTCTCGTGGGCTCGG |
| 702 | CAAGCAGAAGACGGCATACGAGAT CTAGTACG GTCTCGTGGGCTCGG |
| 703 | CAAGCAGAAGACGGCATACGAGAT TTCTGCCT GTCTCGTGGGCTCGG |
| 704 | CAAGCAGAAGACGGCATACGAGAT GCTCAGGA GTCTCGTGGGCTCGG |
| 705 | CAAGCAGAAGACGGCATACGAGAT AGGAGTCC GTCTCGTGGGCTCGG |
| 706 | CAAGCAGAAGACGGCATACGAGAT CATGCCTA GTCTCGTGGGCTCGG |
| 707 | CAAGCAGAAGACGGCATACGAGAT GTAGAGAG GTCTCGTGGGCTCGG |
| 708 | CAAGCAGAAGACGGCATACGAGAT CCTCTCTG GTCTCGTGGGCTCGG |
| 709 | CAAGCAGAAGACGGCATACGAGAT AGCGTAGC GTCTCGTGGGCTCGG |
| 710 | CAAGCAGAAGACGGCATACGAGAT CAGCCTCG GTCTCGTGGGCTCGG |
| 711 | CAAGCAGAAGACGGCATACGAGAT TGCCTCTT GTCTCGTGGGCTCGG |
| 712 | CAAGCAGAAGACGGCATACGAGAT TCCTCTAC GTCTCGTGGGCTCGG |
| 714 | CAAGCAGAAGACGGCATACGAGAT TCATGAGC GTCTCGTGGGCTCGG |
| 715 | CAAGCAGAAGACGGCATACGAGAT CCTGAGAT GTCTCGTGGGCTCGG |
| 716 | CAAGCAGAAGACGGCATACGAGAT TAGCGAGT GTCTCGTGGGCTCGG |
| 718 | CAAGCAGAAGACGGCATACGAGAT GTAGCTCC GTCTCGTGGGCTCGG |
| 719 | CAAGCAGAAGACGGCATACGAGAT TACTACGC GTCTCGTGGGCTCGG |
| 720 | CAAGCAGAAGACGGCATACGAGAT AGGCTCCG GTCTCGTGGGCTCGG |
| 721 | CAAGCAGAAGACGGCATACGAGAT GCAGCGTA GTCTCGTGGGCTCGG |
| 722 | CAAGCAGAAGACGGCATACGAGAT CTGCGCAT GTCTCGTGGGCTCGG |
| 723 | CAAGCAGAAGACGGCATACGAGAT GAGCGCTA GTCTCGTGGGCTCGG |
| 724 | CAAGCAGAAGACGGCATACGAGAT CGCTCAGT GTCTCGTGGGCTCGG |
| 726 | CAAGCAGAAGACGGCATACGAGAT GTCTTAGG GTCTCGTGGGCTCGG |
| 727 | CAAGCAGAAGACGGCATACGAGAT ACTGATCG GTCTCGTGGGCTCGG |
| 728 | CAAGCAGAAGACGGCATACGAGAT TAGCTGCA GTCTCGTGGGCTCGG |
| 729 | CAAGCAGAAGACGGCATACGAGAT GACGTCGA GTCTCGTGGGCTCGG |
| index primers (nextera) N5xx | |
| 501 | AATGATACGGCGACCACCGAGATCTACAC TAGATCGC TCGTCGGCAGCGTC |
| 502 | AATGATACGGCGACCACCGAGATCTACAC CTCTCTAT TCGTCGGCAGCGTC |
| 503 | AATGATACGGCGACCACCGAGATCTACAC TATCCTCT TCGTCGGCAGCGTC |
| 504 | AATGATACGGCGACCACCGAGATCTACAC AGAGTAGA TCGTCGGCAGCGTC |
| 505 | AATGATACGGCGACCACCGAGATCTACAC GTAAGGAG TCGTCGGCAGCGTC |
| 506 | AATGATACGGCGACCACCGAGATCTACAC ACTGCATA TCGTCGGCAGCGTC |
| 507 | AATGATACGGCGACCACCGAGATCTACAC AAGGAGTA TCGTCGGCAGCGTC |
| 508 | AATGATACGGCGACCACCGAGATCTACAC CTAAGCCT TCGTCGGCAGCGTC |
| 510 | AATGATACGGCGACCACCGAGATCTACAC CGTCTAAT TCGTCGGCAGCGTC |
| 511 | AATGATACGGCGACCACCGAGATCTACAC TCTCTCCG TCGTCGGCAGCGTC |
| 513 | AATGATACGGCGACCACCGAGATCTACAC TCGACTAG TCGTCGGCAGCGTC |
| 515 | AATGATACGGCGACCACCGAGATCTACAC TTCTAGCT TCGTCGGCAGCGTC |
| 516 | AATGATACGGCGACCACCGAGATCTACAC CCTAGAGT TCGTCGGCAGCGTC |
| 517 | AATGATACGGCGACCACCGAGATCTACAC GCGTAAGA TCGTCGGCAGCGTC |
| 518 | AATGATACGGCGACCACCGAGATCTACAC CTATTAAG TCGTCGGCAGCGTC |
| 520 | AATGATACGGCGACCACCGAGATCTACAC AAGGCTAT TCGTCGGCAGCGTC |
| 521 | AATGATACGGCGACCACCGAGATCTACAC GAGCCTTA TCGTCGGCAGCGTC |
| 522 | AATGATACGGCGACCACCGAGATCTACAC TTATGCGA TCGTCGGCAGCGTC |
