## Supplementary material for "Student-Driven Microbiome Exploration: A Low-Cost 16S rRNA Sequencing Curriculum for Undergraduate Biology Education": Poster S2

### Evaluating Disinfectant Wipe Impact on Gym Equipment's Surface Microbiome Using 16S rRNA Sequencing

Ana Sofía Vindas, Hassan Barakat, Makenzie Bolton, Jingjing Cheng, Ackshaya Muthu Saravanan, Xiangpeng Li*
Department of Chemistry and Biochemistry, Florida State University,*

#### Introduction

##### Background

Microorganisms are readily transferred through human contact, making shared gym equipment potential reservoirs for bacterial communities. Assessing how these communities respond to disinfection is critical for understanding hygiene practices and reducing potential health risks.

##### Objectives

- Characterize bacterial community composition across gym equipment surface types (dumbbell, bench, yoga mat) using 16S rRNA gene sequencing.
- Evaluate diversity changes after disinfection with branded (Clorox) vs. non-branded (alcohol) wipes.

##### Public Health Relevance

Clinically relevant bacteria such as Staphylococcus, Acinetobacter, and Pseudomonas may persist on gym equipment even after disinfection, posing a potential risk to users.

#### Methods

##### Sample Collection

Swabs from dumbbells, benches, and yoga mats (n = 3 replicates each), before and 10 min after wiping with branded (Clorox) or non-branded (alcohol) wipes.

##### Sequencing

DNA extraction → 16S rRNA V4 amplicon sequencing → QIIME2 (DADA2 denoising, SILVA taxonomy)

##### Statistical Analyses

Alpha diversity: Faith's PD & Shannon — Wilcoxon rank-sum and signed-rank tests
Beta diversity: Weighted/Unweighted UniFrac PCoA — PERMANOVA
Differential abundance: ANCOM-BC (q < 0.05, |LFC| > 1)
Community overlap: Venn diagrams (genus ≥ 0.1% in ≥ 1 sample)

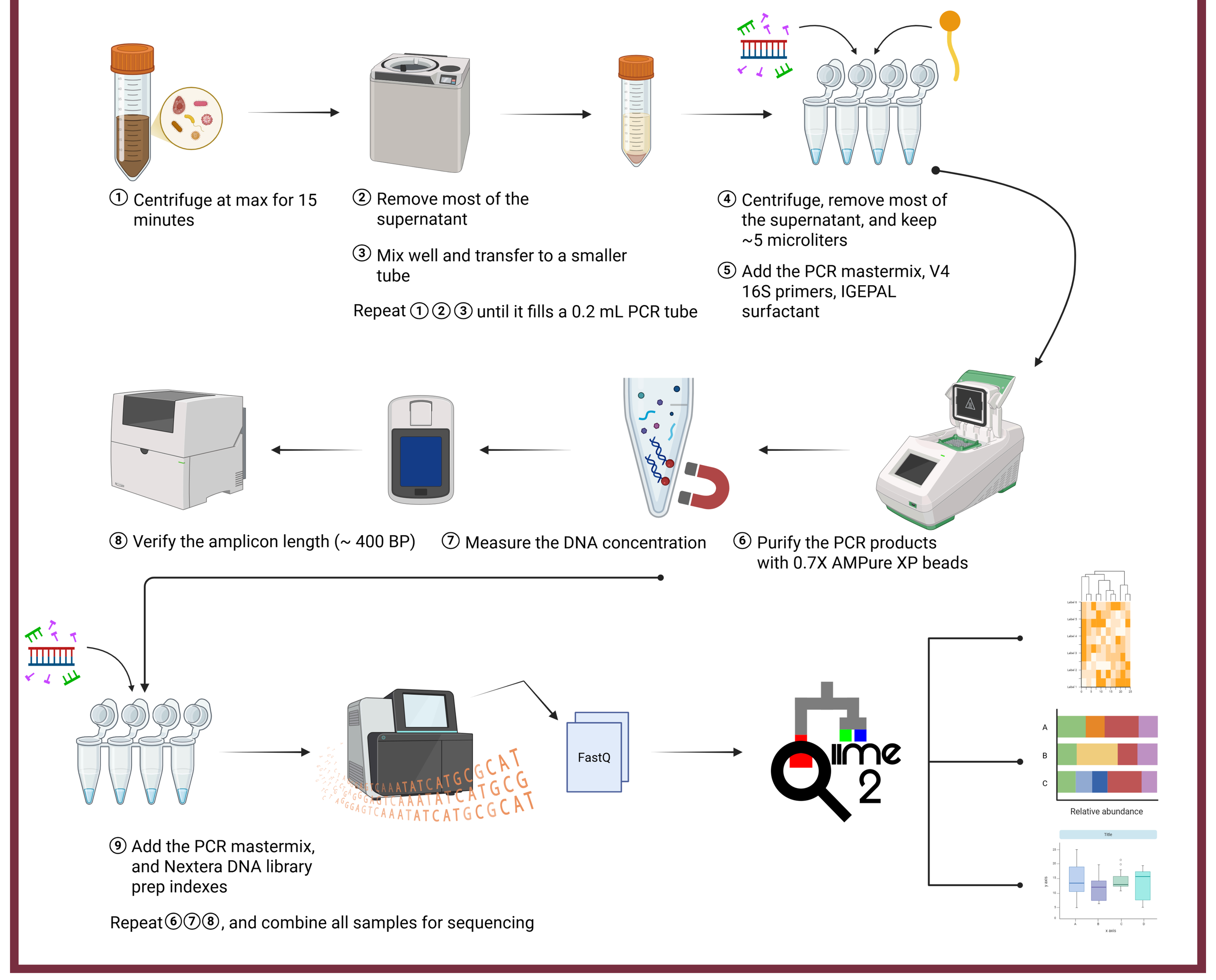


#### Phylum-Level Bacterial Composition Across All Samples

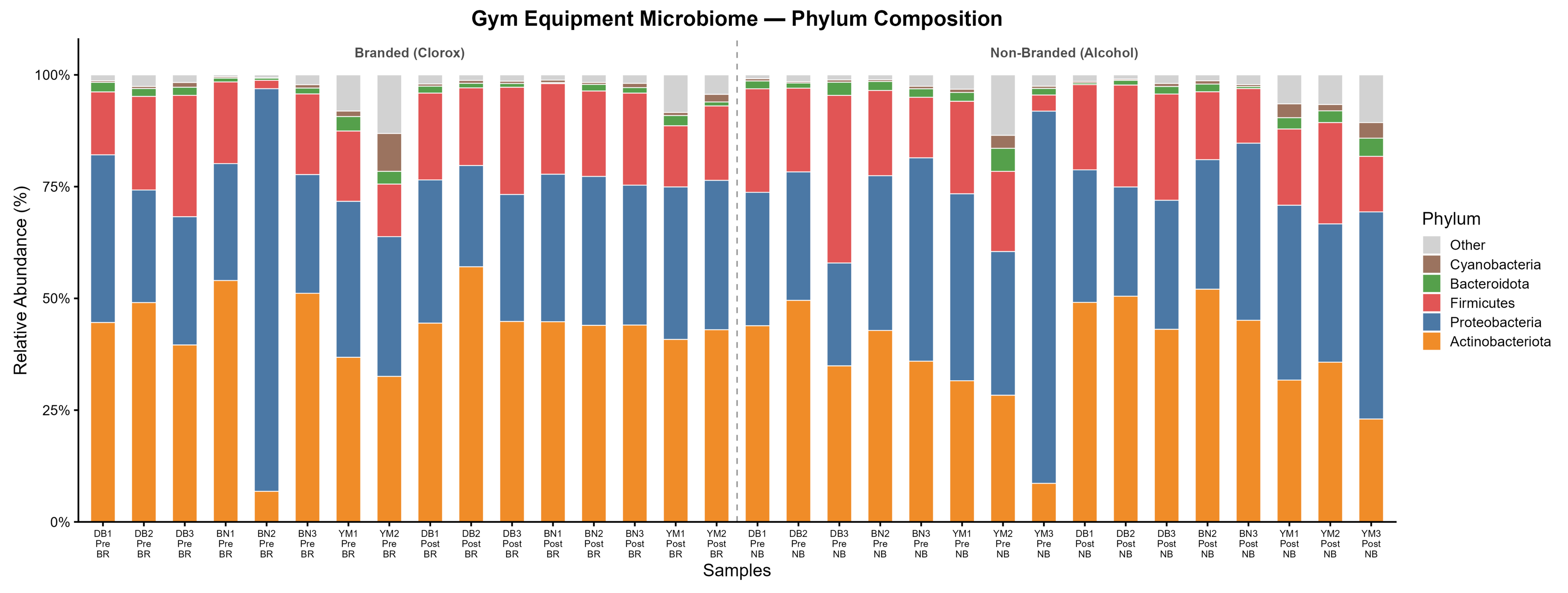


Fig. 1. Relative abundance (%) of bacterial phyla (≥1% mean) across all samples ordered by wipe brand (Branded → Non-Branded), timepoint (Pre → Post), and equipment type. Actinobacteriota, Proteobacteria, and Firmicutes dominate all samples, consistent with a human skin-associated microbiome.

#### Top 50 Genera Heatmap

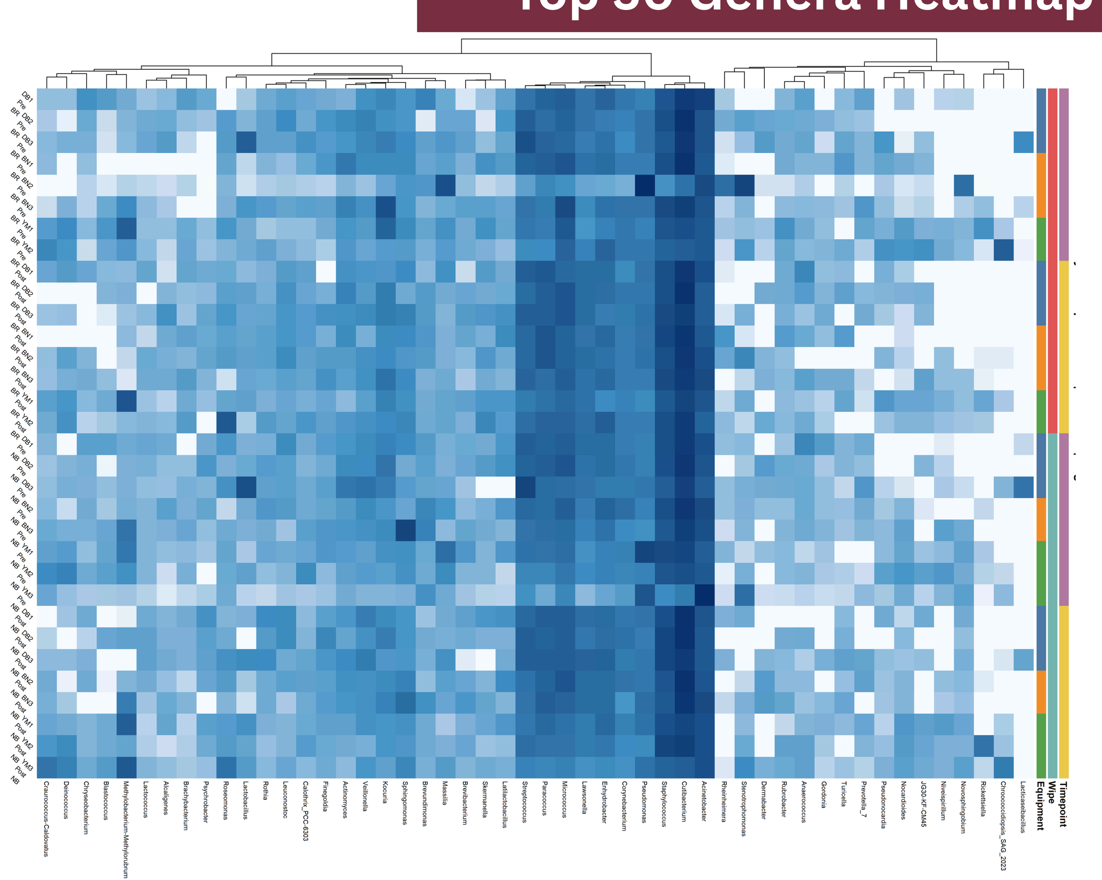


Fig. 5. $Log_{10}$ relative abundance of top 50 genera (Euclidean clustering, complete linkage). Annotation tracks: timepoint, wipe brand, equipment type. Cutibacterium, Acinetobacter, and Staphylococcus form dominant high-abundance clusters across all surfaces.

#### Alpha Diversity by Equipment Type

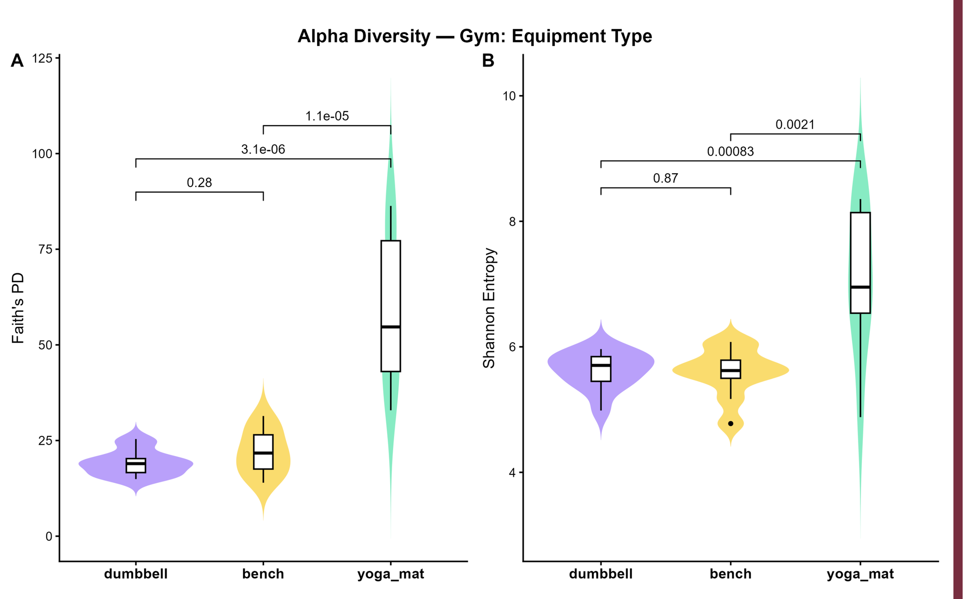


Fig. 2. Faith's PD (A) and Shannon Entropy (B) by equipment type. Yoga mats exhibited significantly higher diversity than dumbbells (Faith's PD: $p = 3.1\times10^{-6}$; Shannon: $p = 8.3\times10^{-4}$) and benches ($p = 1.1\times10^{-5}$; $p = 0.0021$). Wilcoxon rank-sum; n = 12 per group.

#### Shared Genera by Equipment

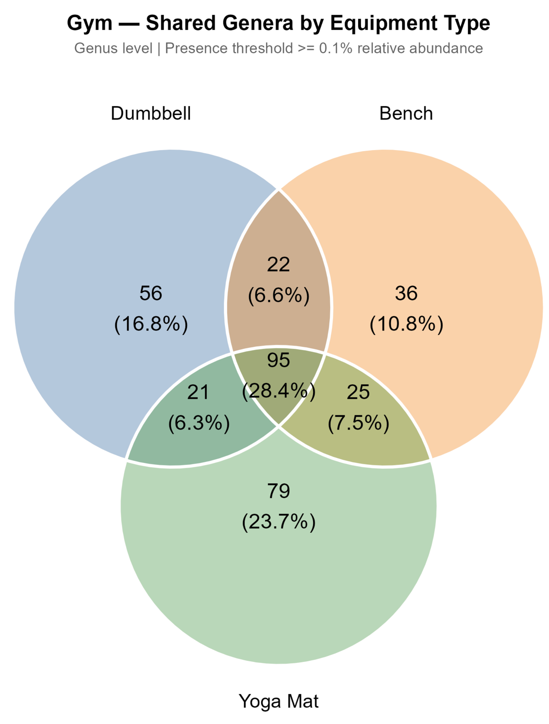


Fig. 3. Venn diagram of genera (≥0.1%) per equipment type. A core of 95 genera (28.4%) was shared across all three surfaces; yoga mats contributed the most unique genera (79; 23.7%), while dumbbells contributed the most among hard surfaces (56; 16.8%).

#### Clinically Relevant Genera Before vs. After Wiping

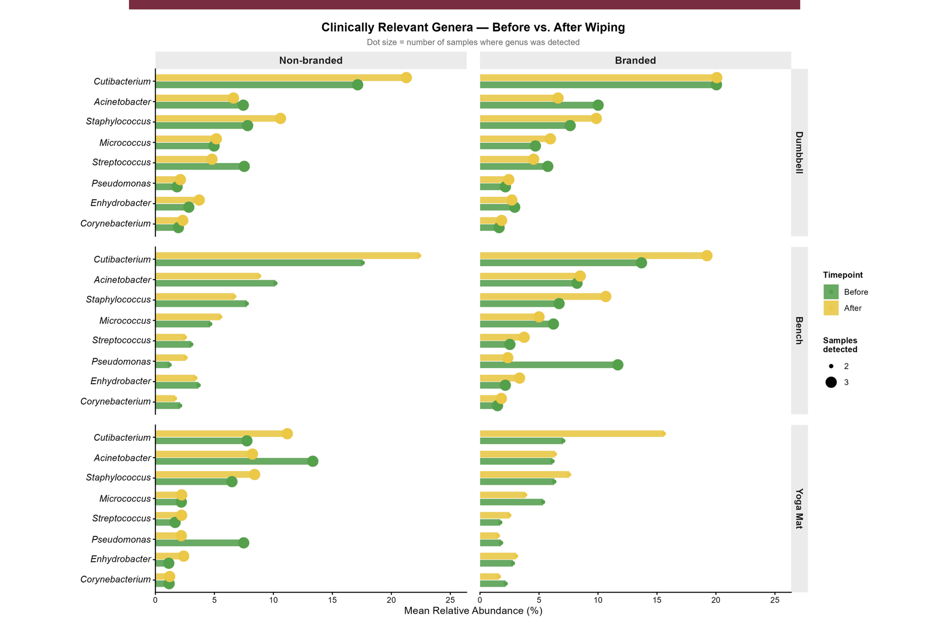


Fig. 6. Mean relative abundance of 8 clinically relevant genera before (green) and after (yellow) wiping, by equipment type × wipe brand. Dot size = samples detected (max n = 3). Cutibacterium dominated across all conditions; Staphylococcus and Acinetobacter remained detectable post-wiping under both wipes

#### Paired Diversity Before vs. After

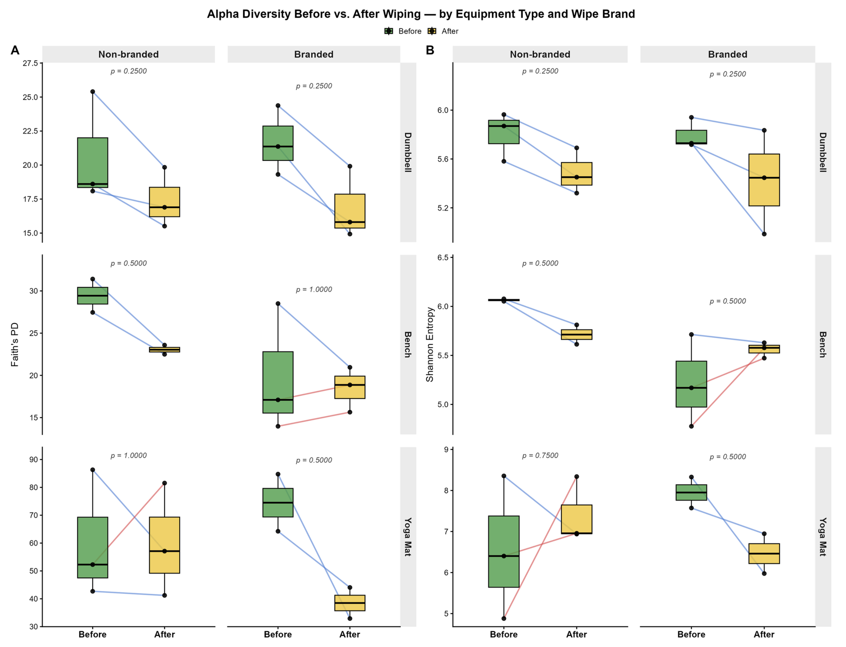


Fig. 7. Faith's PD (A) and Shannon (B) before (green) and after (yellow) wiping per equipment × wipe brand. Blue = decreased; red = increased. Wilcoxon signed-rank; discrete p-values (n = 2–3 pairs). No facet reached significance.

#### Differential Abundance: Before vs. After Wiping

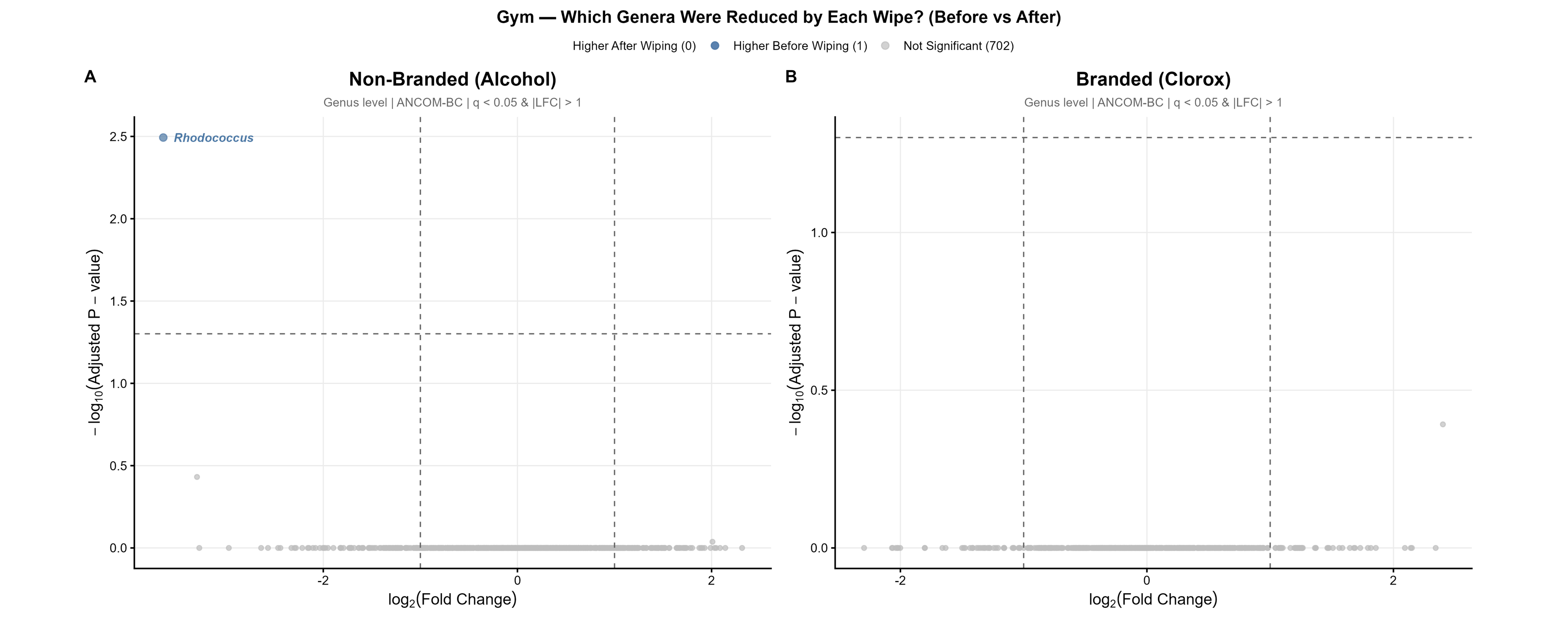


Fig. 4. Volcano plots of ANCOM-BC differential abundance (before vs. after wiping) for non-branded alcohol wipe (A) and branded Clorox wipe (B). Genus-level; q < 0.05 and |LFC| > 1. Only Rhodococcus (blue; higher before wiping) met significance criteria for the non-branded wipe; no taxa were significant for the branded wipe, indicating limited compositional shifts at 10 min post-wiping.

#### Discussion

- Bacterial communities on gym equipment are dominated by skin-associated genera (e.g., Cutibacterium, Staphylococcus, Acinetobacter) and show consistent composition across surface types.
- Yoga mats exhibit higher microbial diversity than hard surfaces, likely due to surface porosity, while dumbbells and benches show similar community profiles.
- Disinfection with either alcohol or Clorox wipes does not significantly reduce diversity or eliminate clinically relevant genera after 10 minutes.
- Branded and non-branded wipes perform similarly, indicating minimal differences in their impact on microbial community structure.

#### Limitations

**Small sample size:** (n = 3 per group), limiting statistical power.
**Single timepoint:** 10 min post-wiping only.
**Amplicon sequencing only:** genus-level; cannot distinguish viable vs. dead cells
**No environmental controls:** air or skin samples not collected.
**One facility**: findings may not generalize.

#### Future Work

→ Increase sample size, n ≥ 10 per group.
→ Multi-timepoint post-wiping (30 min, 1 hr, 24 hr)
→ Incorporate PMA viability assay.
→ Apply shotgun metagenomics for higher resolution & resistome profiling
→ Expand to multiple facilities
→ Compare surface materials (rubber, metal, vinyl, foam

**Acknowledgment**: This work was supported by Florida State University Startup Fund and Florida State University First Year Assistant Professor Award (FYAP).



Li Lab @FSU | DEPARTMENT OF CHEMISTRY & BIOCHEMISTRY
COLLEGE OF ARTS & SCIENCES
