## Supplementary material for "Student-Driven Microbiome Exploration: A Low-Cost 16S rRNA Sequencing Curriculum for Undergraduate Biology Education": Poster S1

### The Microbiology of Celebration: How the Wescott Fountain 21st Birthday Tradition Shapes its Water Microbiome

Kyle Lee, Hassan Barakat, Makenzie Bolton, Jingjing Cheng, Ackshaya Muthu Saravanan, Xiangpeng Li*
Department of Chemistry and Biochemistry, Florida State University, *

#### INTRODUCTION

Florida State University's long lasting Wescott Fountain tradition, where students are tossed into the fountain at midnight to celebrate their 21st birthday, creates a uniquely high contact environment that may influence the microbial communities present in the water. To explore how such frequent physical interactions shape microbial structure, we conducted a two-week sampling study that included a baseline week and a week representing increased student activity. Water was collected two times daily and processed through a validated workflow. Sequencing will allow us to characterize bacterial diversity and detect shifts in community structure between low activity and high activity periods. We expect that samples collected during heightened student interaction will show noticeable changes in microbial composition and diversity compared to baseline conditions, reflecting the impact of human contact on the fountain's water microbiome. This work will establish an initial framework for understanding how student-driven behaviors influence microbial dynamics in public water features on college campuses.

#### WORKFLOW

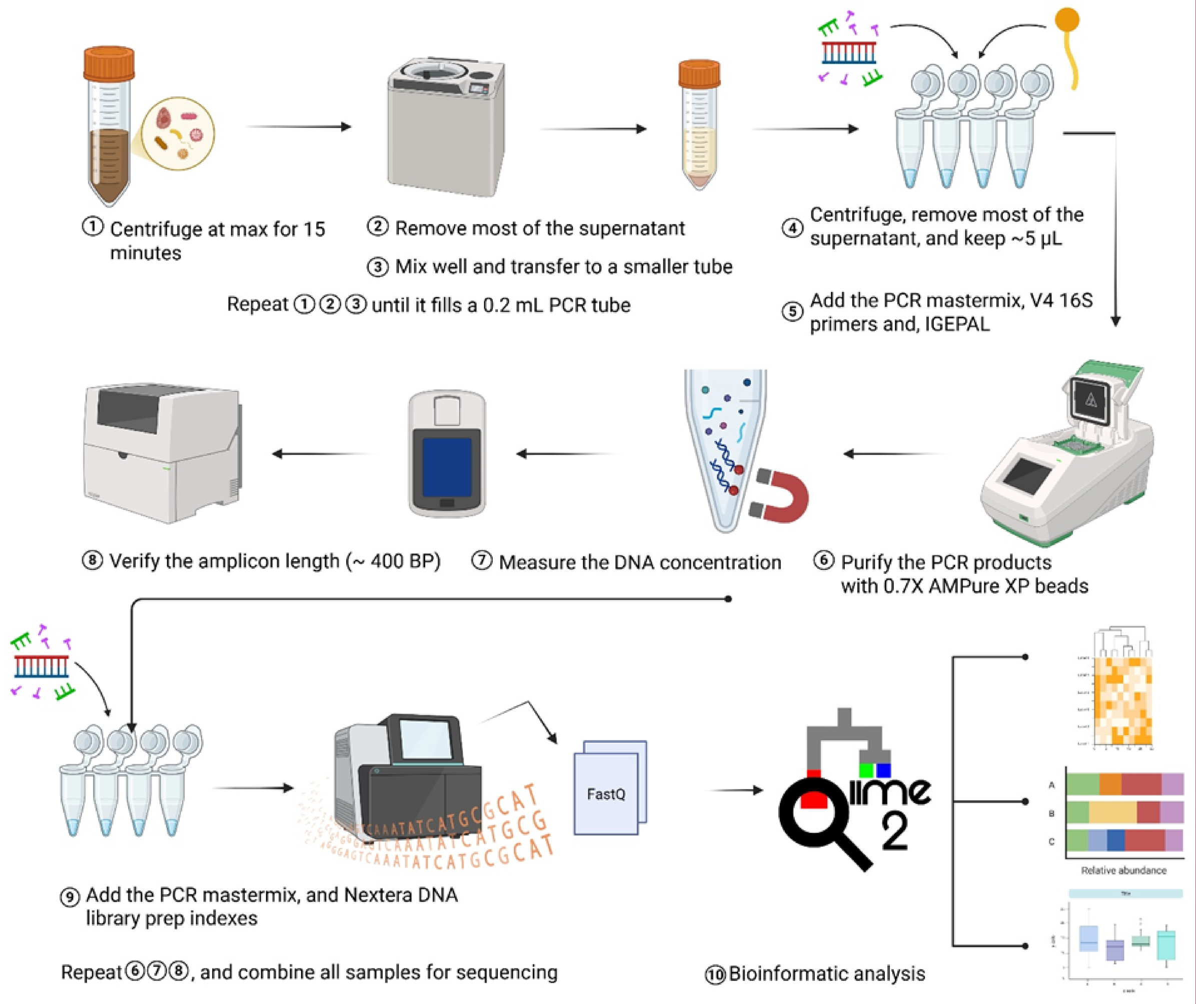


Bulk water samples were collected from the Westcott Fountain at two daily timepoints (afternoon and midnight) across 6 sampling days, capturing three naturally occurring conditions: time of day, rain events, and student activity (birthday fountain-throwing events). Collected samples were processed using IGEPAL, a surfactant that enables DirectPCR without a DNA extraction step, followed by 16S rRNA V4 amplification, sequencing, and bioinformatic analysis using the QIIME2 pipeline.

#### RESULTS

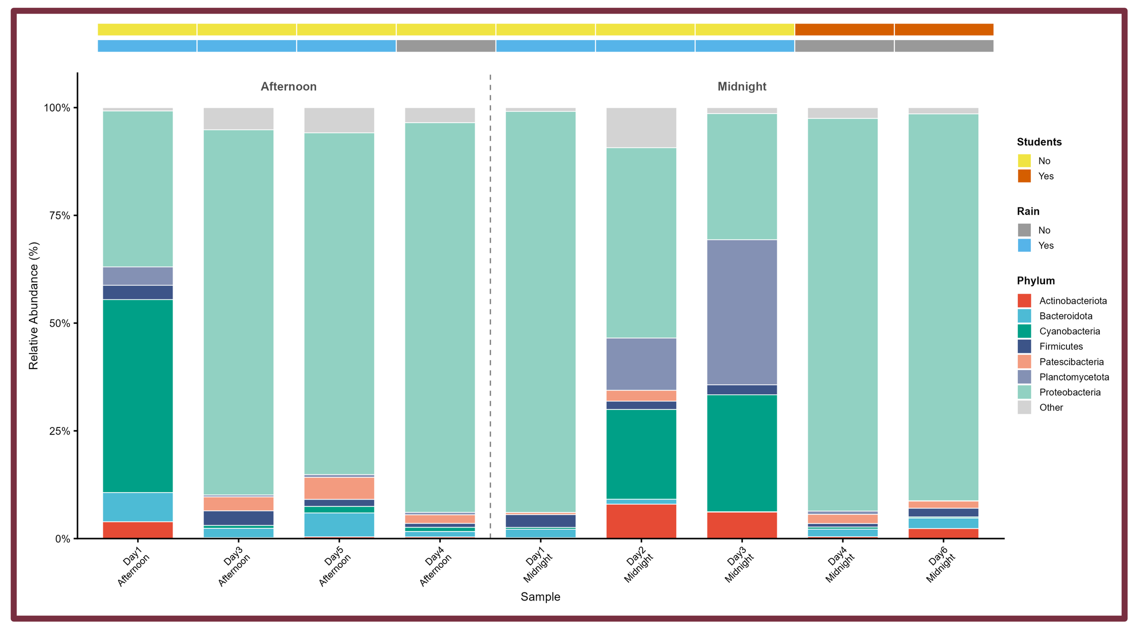

**Fig. 1.** Relative abundance (%) of bacterial phyla across all fountain water samples. Samples are ordered by timepoint (Afternoon → Midnight); dashed line separates groups. Annotation tracks indicate student activity (orange = yes, yellow = no) and rain event (blue = yes, gray = no). Cyanobacteria and Proteobacteria dominate most samples, with notable Planctomycetota enrichment on Day 3 Midnight coinciding with a rain event.

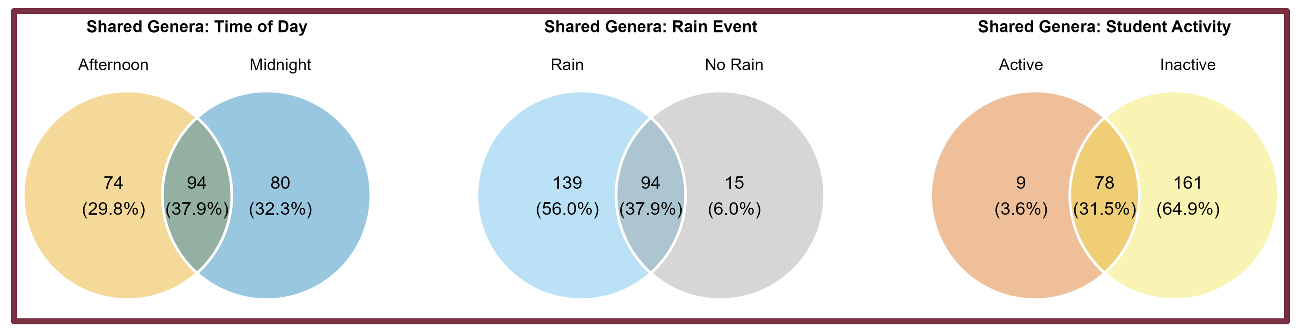


**Fig. 5.** Venn diagrams of bacterial genera (≥ 0.1% in ≥ 1 sample) shared across groups for three variables: time of day (left), rain event (center), and student activity (right). A core of 94 genera (37.9%) was shared across time of day and rain groups. The student activity comparison was highly asymmetric: 161 genera (64.9%) were exclusive to inactive periods vs. only 9 (3.6%) to active periods, likely reflecting the larger number of inactive samples. Compositional enrichment during activity is better captured in Fig. 3.

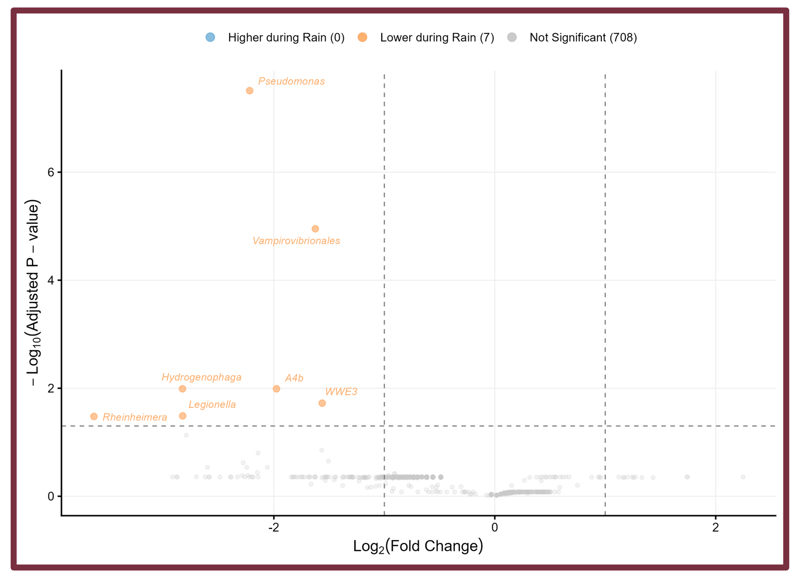

**Fig. 4.** ANCOM-BC differential abundance analysis comparing rain vs. no rain conditions. Genus level; q < 0.05 and |LFC| > 1. Seven genera were significantly lower during rain (orange) including *Pseudomonas*, *Legionella*, and *Vampirovibrionales*, with none significantly higher, consistent with a rainfall dilution effect on the resident microbial community.

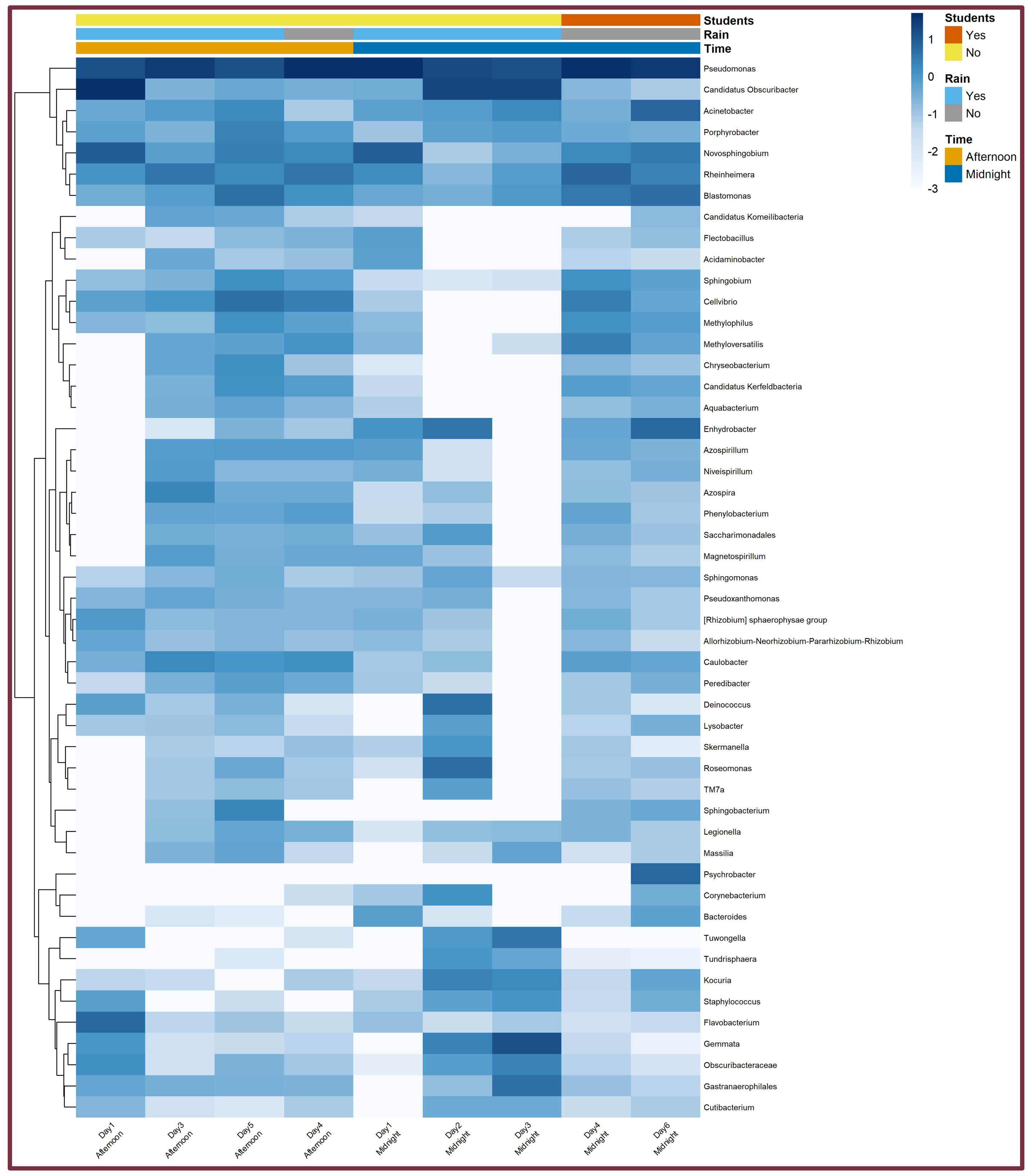

**Fig. 2.** Log₁₀ relative abundance heatmap of top bacterial genera across all samples (Euclidean clustering, complete linkage). Column annotation tracks indicate student activity, rain event, and time of day. A pseudocount of 0.001 was added prior to log-transformation. *Pseudomonas*, *Acinetobacter*, and *Candidatus Obscuribacter* were the most consistently dominant genera, with pronounced abundance shifts associated with student activity and rain events.

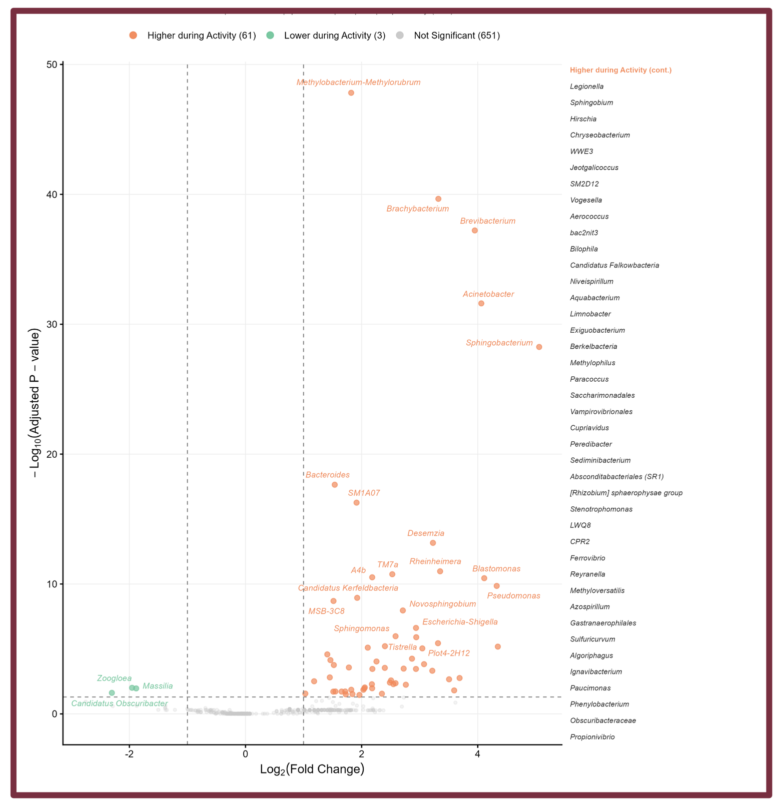

**Fig. 3.** ANCOM-BC differential abundance analysis comparing student activity vs. no activity (exploratory; n = 2 active samples). Genus level; q < 0.05 and |LFC| > 1. 61 genera were significantly higher during activity (orange) including *Brevibacterium*, *Brachybacterium*, *Acinetobacter*, and *Escherichia-Shigella*, consistent with direct human microbiome input during immersion. Only 3 genera were lower during activity (green). Results should be interpreted cautiously given the small active sample size.

#### DISCUSSION, CONCLUSION & FUTURE DIRECTIONS

Student birthday fountain-throwing events drove the most pronounced microbial shift observed, with 61 genera significantly enriched during active periods spanning skin (*Brevibacterium*, *Brachybacterium*, *Acinetobacter*), gut (*Bacteroides*, *Escherichia-Shigella*), and oral-associated taxa which is consistent with direct human microbiome deposition during full-body immersion. Rain events had the opposite effect: 7 resident genera including *Pseudomonas* and *Legionella* were significantly reduced during rainfall, consistent with a dilution of the resident community rather than introduction of new taxa. Time of day alone showed no detectable effect on community composition (0 significant taxa, Midnight vs. Afternoon), suggesting diurnal cycling is not a primary driver of microbial variation in this system. Together, these findings indicate that the Westcott fountain microbiome is shaped primarily by episodic perturbations, human immersion and rainfall, rather than stable temporal cycling. The student activity comparison is limited by small sample size (n = 2 active samples) and should be considered exploratory. **In conclusion**, student fountain-throwing significantly enriches human-associated taxa in fountain water; rain events dilute the resident microbiome including *Legionella*; and time of day has no independent effect on community composition. **Future work** should increase active event sampling (n ≥ 10) with paired pre/post collection to quantify perturbation magnitude and recovery time, extend sampling across seasons, include water chemistry covariates (pH, chlorine, turbidity), apply shotgun metagenomics for species-level resolution and resistome profiling, and compare Westcott to other campus water features to contextualize baseline community composition.

#### ACKNOWLEDGEMENTS

This work was supported by Florida State University Startup Fund and Florida State University First Year Assistant Professor Award (FYAP).

*Know more about our lab: Li-lab.us*
